# How does the mood stabilizer lithium bind ATP, the energy currency of the cell

**DOI:** 10.1101/637900

**Authors:** A. Haimovich, A. Goldbourt

## Abstract

Lithium, in the form of a salt, is a mood stabilizer and a leading drug for the treatment of bipolar disorder. It has a very narrow therapeutic range and a variety of side effects. Lithium can replace magnesium and other cations in enzymes and small molecules, among them ATP, thereby affecting and inhibiting many biochemical pathways. The form of binding of lithium ions to ATP is not known.

Here we extract the binding environment of lithium in solid ATP using a multi-nuclear multi-dimensional solid-state NMR approach.

We determine that the coordination sphere of lithium includes, at a distance of 3.0(±0.4) Å, three phosphates; the two phosphates closest to the ribose ring from one ATP molecule, and the middle phosphate from another ATP molecule. A water molecule most probably completes the fourth coordination. Despite the use of excess lithium in the preparations, sodium ions still remain bound to the sample, at distances of 4.3-5.5 Å from Li, and coordinate the first phosphate and two terminal phosphates.

In conclusion**, s**olid-state NMR enables to unravel the exact coordination of lithium in ATP showing binding to three phosphates from two molecules, none of which are the terminal gamma phosphate. The methods we use are applicable to study lithium bound to a variety of ATP-bound enzymes, or to other cellular targets of lithium, consequently suggesting a molecular basis for its mode of action.

## 1. Introduction

The adenosine triphosphate (ATP) molecule is of paramount importance, enabling myriad processes including energy transport within cells. ATP is composed of a purine base called adenine, a ribose, and three phosphate groups. Hydrolysis of ATP to ADP leaving a free inorganic phosphate (Pi) unit is energetically favorable, having a large negative free energy. It facilitates many biochemical processes by coupling to such reactions, thus reducing their energy requirements. Such processes may or may not involve the direct transfer of the Pi unit. ATP takes part in many cellular processes and signaling pathways - a simple example being the first stage of glycolysis where the Pi group from ATP is attached to glucose generating glucose-6-phospahtes, G6P. ATP also provides energy to allow mechanical motions such as the movement of myosin on actin filaments and serves as an allosteric modulator to enzymes.

ATP has been linked with the pathogenesis of several diseases including type 2 diabetes, Alzheimer’s disease and bipolar disorder.[1,2] It binds to a variety of metal ions, and its metal bound form is the substrate for numerous enzymes. Physiologically, Mg^2+^ is the most relevant metal ion that binds ATP.

Lithium has many biological molecular targets, including molecules that are neurotrophic and molecules related with important signal transduction pathways.[3,4] It is an inhibitor of phosphatases (e.g. inositol monophosphatase) and kinases, and it interacts with phosphorylated substrates such as ATP. In its magnesium bound form ATP is a biological substrate for GSK-3β (glycogen synthase kinase-3β) and thus lithium causes its inhibition. Lithium enters the cells via voltage-sensitive sodium channels and can affect intracellular sodium and calcium levels.[5]

The structure and binding of metal complexes of ATP has been studied extensively, and the association constants of different metals to ATP were measured by various methods to some diversity.[6–15] These solution state studies report on a 1:1 ionic complex formation, dominated by MgATP^2-^ species. However, the existence of a neutral Mg_2_ATP complex was also observed[9,13] and the possible formation of the complex, Mg(ATP)_2_^6-^ was suggested.[8] Measurements have shown that for the MATP^2-^ complex (M≡Metal), among divalent cations Mg^2+^ is a strong, but not the strongest, binder,[6,7,9,11,12]. Among monovalent cations such as Li^+^, Na^+^, and K^+^, Li^+^ has the strongest binding affinity. It was also determined that monovalent cations compete with magnesium binding to ATP.[11]

The ATP molecule has been studied by various NMR techniques. Metal binding has been shown to affect ^31^P chemical shifts and ^31^P-^31^P scalar couplings in solution.[13–21] Ramirez et al. suggested that changes in the chemical shifts resulted from conformational changes of the phosphate chain and were not necessarily indicative of metal binding to a specific phosphate group.[18] On the other hand, Haake et al. concluded that changes in ^31^P chemical shifts must only be due to direct association with the metal.[14] Solid state Magic-angle spinning (MAS) ^31^P NMR studies of disodium-ATP[21,22] related changes in the phosphorous spectra also to the hydration state.

Sodium ATP structural crystallographic studies[23–25] in the solid state revealed two ATP molecules in the asymmetric unit, with four chemically inequivalent sodium sites and six water molecules. Two of the sodium sites form a dimer coordinating to the phosphate oxygens and a base nitrogen of both ATP molecules. The other two link the ATP dimers in the extended structure. The phosphate chains have a folded conformation and form partial helices; molecule A forms a left handed helix and molecule B forms a right handed helix (see figure #16 of [23]). The ribose rings also have different conformations; molecule A has an N-type sugar pucker while molecule B has an S-type sugar pucker. The adenine bases are stacked almost parallel to each other. The existence of four inequivalent sodium sites in sodium ATP has also been identified in ^23^Na multiple-quantum MAS (MQMAS) NMR studies combined with quantum mechanical calculation methods.[26–29]

Magnesium ATP could only be crystalized with bis(2-pyridyl)amine (bipyam) creating [Mg(H_2_O)_6_][Hbipyam]_2_[Mg(HATP)_2_]·nH_2_O complexes.[30–32] Mg^2+^ occupies two inequivalent sites (see Figure S9 in the SI), where one Mg^2+^ cation forms an ATP-metal complex and the other binds six water molecules. The ATP-metal complex has an octahedral geometry with the metal bound to all phosphate groups, three from each ATP molecule. No bonding interactions were observed with the purine base or the ribose. ^25^Mg MQMAS NMR studies also revealed the existence of two magnesium sites.[33] The phosphate chains in MgATP have a folded conformation, similarly to Na_2_ATP, and are part of a network of hydrogen bonds formed with the metal-water complex, the adenine nitrogens, the ribose hydroxy oxygens and free water. Both ribose rings show an S-type sugar pucker conformation as the asymmetric unit contains a single MgATP moiety. The purine base has strong stacking interactions with the bipyam molecules and not with other bases. The metal-water complex was suggested to play a role in phosphoryl transfer due to its coordination with the phosphates chain.

Evidence for competition in solution between Li^+^ and Mg^2+^ for binding sites in ATP was obtained using fluorescence, Isothermal titration calorimetry (ITC), spin-lattice ^7^Li T_1_ relaxation time measurements and ^31^P NMR.[11,15,20,34] It was argued that binding of the metals to the base or sugar moieties was negligible and that Li^+^ and Mg^2+^ compete for phosphate binding sites. Furthermore it was shown that magnesium affinity is greater than that of lithium and that lithium binding constants to the phosphate group were somewhat greater in the presence of magnesium. While both LiATP and Li_2_ATP species exist, at pH>7 the Li_2_ATP species predominates. Both Mg^2+^ and Li^+^ were said to bind to all three phosphate groups. A recent ^7^Li relaxation study in solution suggested that lithium binds MgATP to form a physiologically relevant Mg-Li-ATP complex, where Mg^2+^ coordinates to P_β_, P_γ_, and water molecules, and Li^+^ coordinates to P_γ_ and water molecules (one bridging Mg^2+^).[35] Free energy calculations on Mg-Li-ATP complexes concluded that Mg^2+^ binds all phosphates, Li^+^ binds P_β_ and P_γ_, and the two metals are linked by a hydroxide bridge. It was also determined that lithium binding to MgATP did not significantly alter the complex conformation implicating on the biological action of lithium.[36]

Here we utilize a multi-nuclear MAS ssNMR approach to study a lithium-ATP-water complex, prepared from sodium-ATP, in order to find the lithium binding site. We show that there are two inequivalent molecules in the unit cell, and we suggest a model for lithium binding that involves P_α_ and P_β_ of one ATP molecule, and P_β’_ of the second ATP molecule. Excess lithium does not replace all sodium ions in the complex, which bind both P_γ_ phosphates.

## 2. Materials and Methods

### 2.1. Preparation of a lithium-ATP-water complex

Li-ATP was prepared by a small modification of the diffusion procedure described by Potrzebowski et al.[22] A 20:1 mole ratio of LiCl and Na_2_ATP·3H_2_O (purchased from Fisher Bioreagents) was dissolved in purified water and isopropyl alcohol (IPA) was added to a ratio of 1:2.5 v/v Water:IPA. The flask containing the solution was then placed in a vessel filled with pure IPA and was left overnight at room temperature to crystalize. The solid material was filtered out, washed with IPA, and dried on a Büchner funnel. In this manner three ATP sample types were prepared that differ in the amount of lithium-7: (i) {23-24%-^7^Li}-Li-ATP: a mixture of ∼25 w/w% natural abundance LiCl and ∼75 w/w% ^6^LiCl; (ii) {36%-Li}-Li-ATP; (iii) {92%-Li}-Li-ATP termed NA Li-ATP – this is the natural abundance sample.

A trihydrate sodium-ATP-water complex was recrystallized following the diffusion procedure described previously.[22] Prior to recrystallization the sample showed broad ^31^P lines indicating lack of a clear crystalline state, or a mixture of hydration states.

### 2.2. NMR Instrumentation and Referencing

The majority of solid state NMR experiments were performed on a Bruker Avance-III spectrometer operating at a magnetic field of 14.1 T, corresponding to Larmor frequencies of 600.2 MHz for ^1^H, 242.9 MHz for ^31^P, 233.2 MHz for ^7^Li, and 158.7 MHz for ^23^Na. Experiments were performed using a wide-bore 4 mm probe operating in double resonance ^1^H-^31^P and ^1^H-^7^Li modes, in a ^1^H-^31^P-^7^Li mode using a REDOR BOX (©NMR service GmbH) frequency splitter[37,38], and in triple-resonance ^1^H-^31^P-^23^Na and ^1^H-^7^Li-^23^Na modes. Additional experiments were carried out using a fast-spinning 1.3mm probe operating in double-resonance ^1^H-^31^P or ^1^H-^7^Li modes.

Some measurements were performed on a Bruker Avance-III spectrometer operating at a magnetic field of 9.4 T, corresponding to Larmor frequencies of 400.2 MHz for ^1^H, 162.0 MHz for ^31^P, and 105.9 MHz for ^23^Na using a wide-bore 4mm probe operating in ^1^H-^31^P and ^1^H-^23^Na double-resonance modes.

Spectra were externally referenced to adamantane at 1.8 ppm for ^1^H, to O-phospho-L-Serine at 0.3 ppm for ^31^P, to 1M LiCl solution at 0 ppm for ^7^Li, and to 0.1M NaCl solution at 0.29 ppm for ^23^Na (calibrated by secondary external referencing with adamantane). The experiments were performed at different temperatures and the reported values are the temperatures set at the entrance to the probe, which is lower than the actual sample temperature due to frictional heating during sample spinning. Detailed experimental parameters are given in the captions and in the supplementary information.

### 2.3. Terminology

MgATP and Na_2_ATP have fixed stoichiometries and are termed accordingly. The lithium complex is termed Li-ATP, or Li(Na)ATP since it is mixed with Na^+^ ions with ratios that are not yet determined. We refer to the Li+ and Mg^2+^ ions as lithium and magnesium respectively throughout the text.

### 2.4. Simulations and fitting of distance measurement experiments

All simulations were performed using the software SIMPSON[39]. ^31^P-detected REDOR simulations were carried out for an isolated ^7^Li–^31^P spin pair, and for three (P_2_-Li) and four (P_3_-Li) spin systems taking into account isotropic shifts, CSA, C_Q_ and all homonuclear and heteronuclear interactions. The carrier frequency was set on-resonance for lithium and to the P_α’_ atom for phosphorus. The values of C_Q_ (50 kHz) and of the ^31^P CSA were set according to the experimentally fitted lineshapes in a {36%-Li}-Li-ATP sample. ^31^P-^31^P homonuclear dipolar interactions, and Euler angles for the CSA tensors and the homonuclear dipolar tensors for the first ATP molecule were all adapted from sodium ATP studies.[22] Euler angles for CSA tensors for the second ATP molecule (two ATP molecules exist in the unit cell) were modified to create a semi mirror image. For the analysis of Li-ATP data, simulations were performed for different heteronuclear dipolar coupling constants. ^31^P and ^7^Li detected PM-RESPDOR and ^23^Na detected REDOR simulations were carried out for two (P-Na or Li-Na) or three (P-Na_2_ or Li-Na_2_) spin systems. Parameters for lithium and phosphorous were similar to the above REDOR simulations. For sodium C_Q_ values of 0.76 MHz and 2.0 MHz were used.

## 3. Results & Discussion

### 3.1. Effects of lithium binding on the triphosphate chain

The ^31^P CPMAS spectrum of the Li-ATP complex, prepared according to a protocol which produced a trihydrate Na_2_ATP, is shown in **Figure 1**. The six ^31^P signals belong to two ATP molecules occupying the asymmetric unit, as observed before for sodium ATP. Also, the spectral features are in agreement with the trihydrate form observed for sodium ATP[21,22].

**Figure 1:**
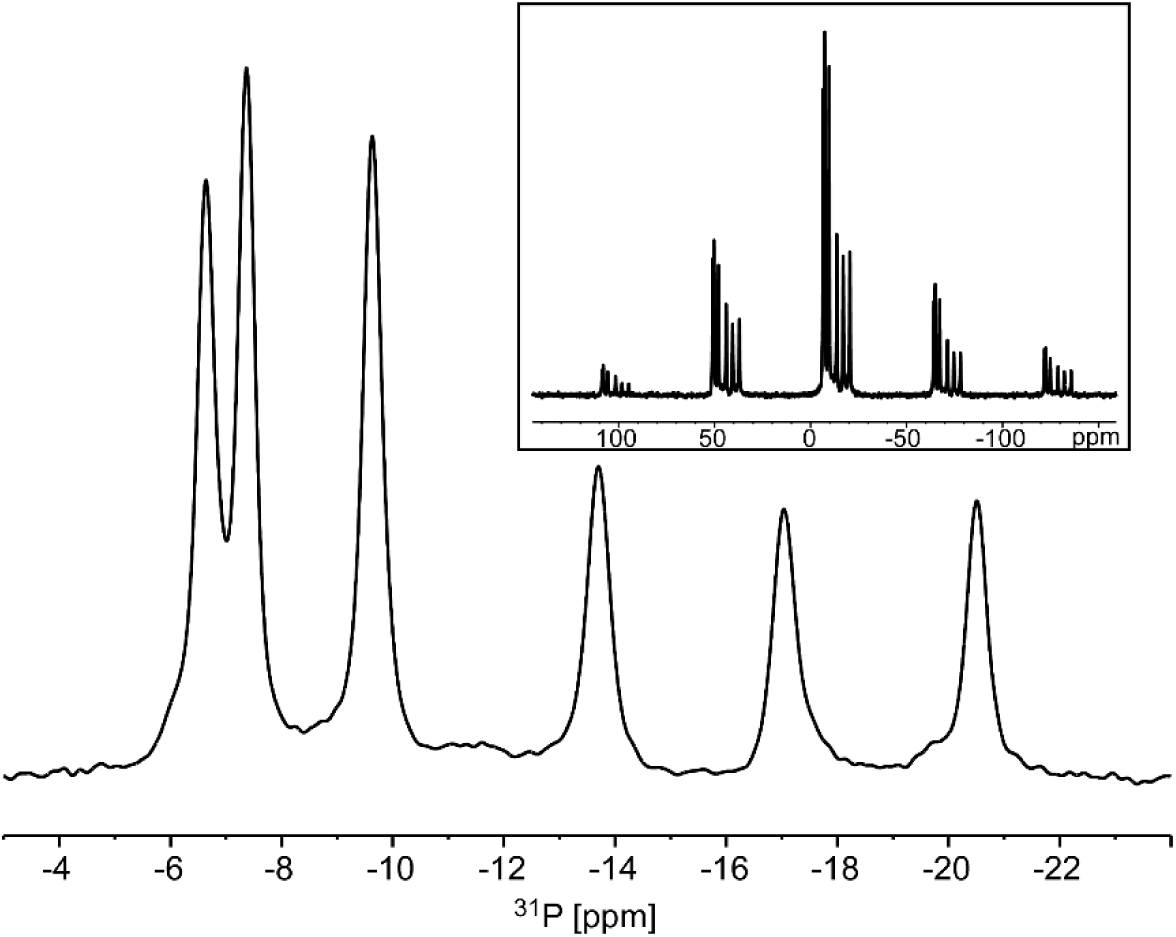
^1^H^−31^P CPMAS center-band spectrum of Li-ATP acquired at a field of 14.1 T and a spinning frequency of 14 kHz. The full spectrum is shown in the inset.

In order to assess the effect of lithium binding on the spectrum, the signals need to be assigned to particular phosphate atoms and in particular to the individual ATP molecules within the dimer. We therefore performed ^31^P-^31^P homonuclear DARR (dipolar assisted rotational resonance[40]) correlation experiments using two mixing times. A DARR experiment conducted with a mixing time of 15 ms (DARR15) is shown in **Figure 2a**. It mainly establishes connectivities within each individual ATP molecule. This approach is similar to the one used to characterize Na_2_ATP[22]. With this spectrum it was possible to identify each molecule unequivocally. By comparison to prior studies (both solid and in solution), we know that γ-phosphates are located at a low-field (high-frequency) position, followed by the α-phosphates and then by the β-phosphates. The locations of the phosphates within the molecule are shown in **Scheme 1.** The relatively strong γ-α correlation suggests that also here, as in the case of sodium and magnesium ATP, the phosphate moiety adopts a bent conformation. Moreover, it is also possible to observe the inter-molecular correlations P_α_-P_γ’_ (in asterisks), while other inter-molecular correlations were significantly weaker. This cross-peak already suggests a form of packing of the two inequivalent molecules.

**Figure 2.**
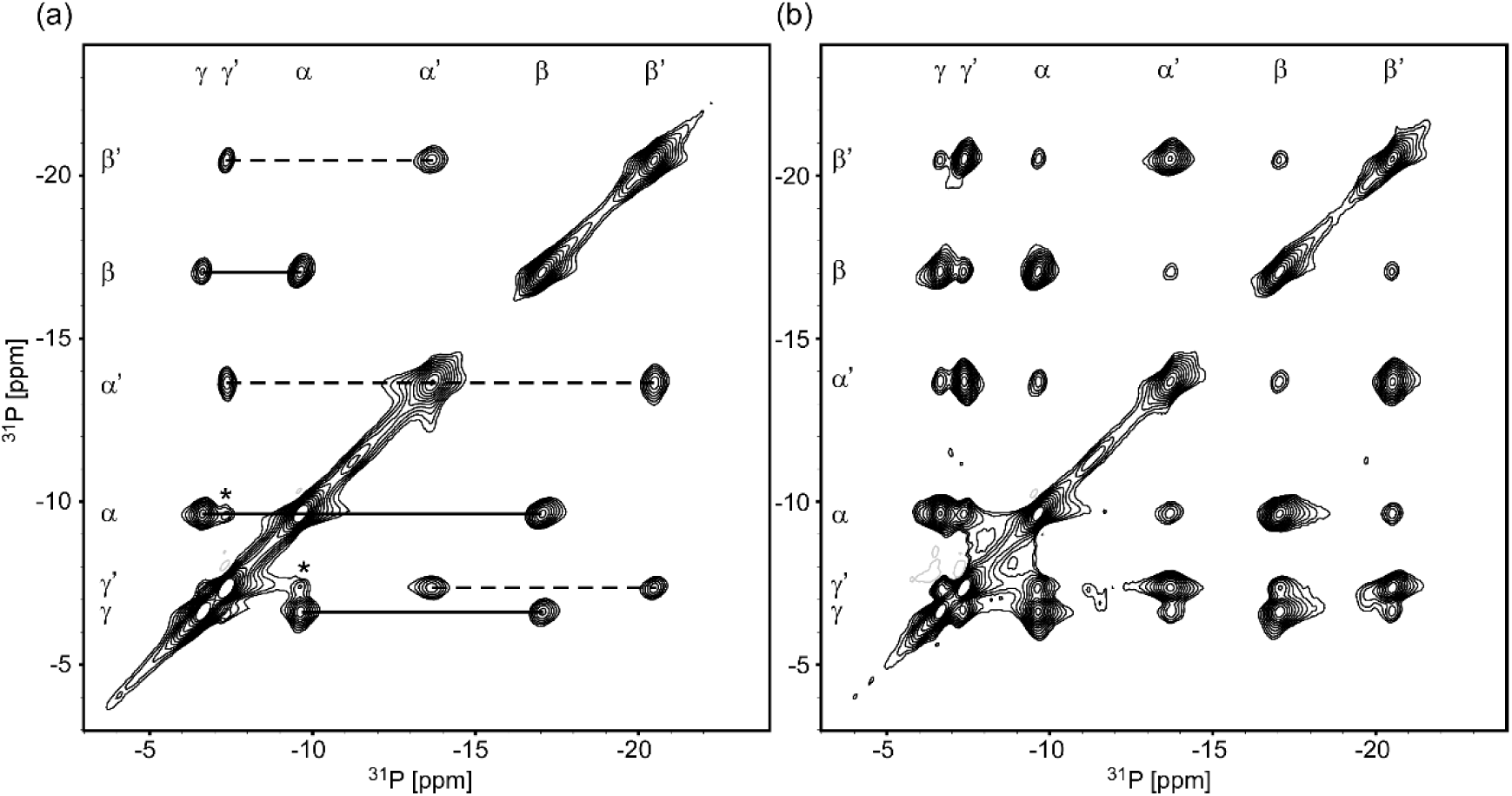
(a) 2D ^31^P-^31^P DARR of Li-ATP acquired with a short mixing time of 15 ms. The lines indicate correlations between adjacent phosphate groups, solid lines correlating signals within phosphates of one ATP molecule (α, β, γ), and the dashed lines for the second ATP molecule (α’, β’, γ’). The asterisks indicate P_α_-P_γ’_ correlations between the two ATP molecules. (b) 2D ^31^P-^31^P DARR of Li-ATP acquired with a mixing time of 100 ms. Many inter-molecular correlations between the two ATP molecules can be observed.

Following the DARR15, CPMAS and prior reports, we could assign all ^31^P signals and fit their CSA values, all of which are given in Table 1.

**Table 1.** Chemical shifts, CSA and asymmetry values for ^31^P in Li-ATP:

| $^{31}\text{P}$ | $P_{\alpha}$ | $P_{\beta}$ | $P_{\gamma}$ | $P_{\alpha'}$ | $P_{\beta'}$ | $P_{\gamma'}$ |
| --- | --- | --- | --- | --- | --- | --- |
| $\delta_{\text{iso}}$ [ppm] | -9.6 | -17.0 | -6.6 | -13.7 | -20.5 | -7.4 |
| $\delta_{\text{aniso}}$ [ppm] | -106.3 | -121.7 | -117.0 | -125.0 | -121.3 | -110.8 |
| $\eta$ | 0.72 | 0.49 | 0.57 | 0.54 | 0.43 | 0.72 |

In order to estimate the organization of the dimer, more inter-molecular correlations should be obtained, and we performed another ^31^P-^31^P DARR experiment using a longer mixing time of 100 ms, allowing longer-range transfers, as shown in **Figure 2b**. From a qualitative analysis of the spectrum we can estimate the relative proximities of the phosphate groups from different molecules. The P_α_-P_γ’_ and P_γ_-P_γ’_ crosspeaks are the strongest of the inter-molecular correlations, and the fact that the P_α_-P_γ’_ are also strongly observed in DARR15 indicates the juxtaposition of these two phosphate groups. Clearly the weakest signals are those connecting P_α_-P_α’_, P_β_-P_β’_, P_α_-P_β’_ and P_α’_-P_β_. Correlations with intermediate strength are observed for P_β_-P_γ’_. Since in the CP spectrum the signal intensities for P_α’_, P_β_ and P_β’_ are about half of the signals of P_α_, P_γ_ and P_γ’_, we also deconvoluted the 2D spectrum (using the software DMFIT[41]), and normalized with respect to the 1D spectral intensities. Similar trends of the cross-peak intensities remain thus strengthening the conclusion that P_α_ and P_γ_ of one molecule are close to P_γ’_ of a second molecule in the unit cell.

Irrespective of lithium position within ATP, our observations suggest a unique organization of the dimer, different from that observed for Na_2_ATP•3H_2_O[22,24], and different from that of MgATP co-crystalized with Bis(2-pyridyl)amine[30]. In the NMR study of Na_2_ATP, double-quantum build-up curves showed close contacts between P_γ_ and P_γ’_, in agreement with the crystallographic short distance of 4.12Å, shorter then intra-molecular P_α_-P_γ_ contacts (see Figure S9 of the SI). No other long-range contacts have been reported, as they are distanced at 6-8Å. Our data reports the shortest contact to be between P_α_ and P_γ’_, followed by P_γ_-P_γ’_ and P_γ_-P_α’_. The equivalent of the P_α_-P_γ’_ contact in MgATP is 6.5Å, and other inter-molecular contacts are ∼5.8Å to P_β_.

**Scheme 1.**
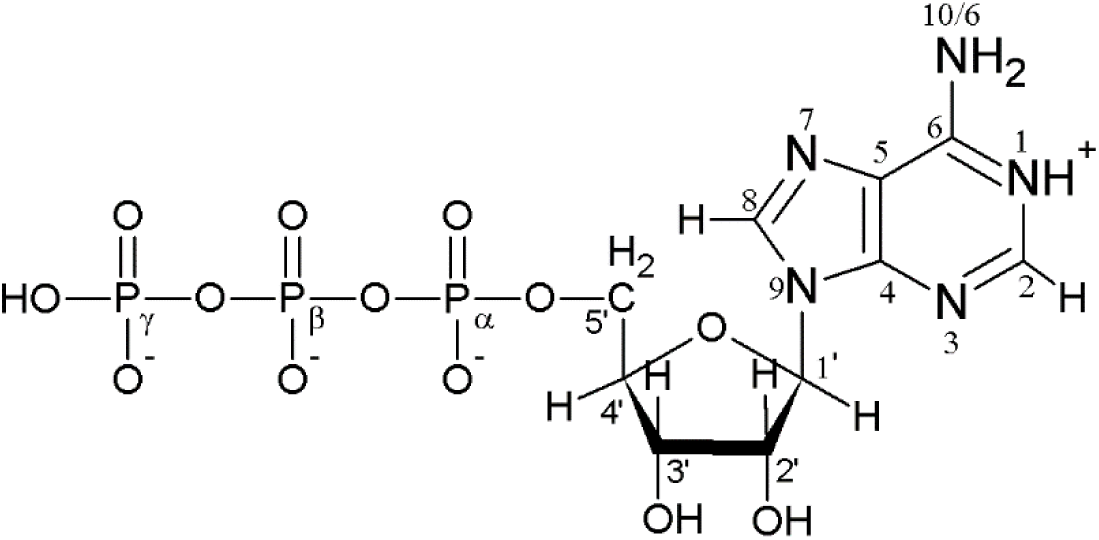
Adenosine triphosphate. Amine base atoms are labeled 1-10, the ribose carbons are labeled 1’-5’ and the phosphate atoms are labeled α, β, γ. The protonation scheme corresponds to an ATP molecule produced at a pH of ∼3. The nomenclature follows that of [23–25]. N10 is labeled N6 in other instances, including the protein data bank. The formal ATP entry in the BMRB (e.g. entry bmse000006) has a different numbering scheme, and is given in the SI.

In order to observe the effect of lithium on ^31^P shifts upon replacement of sodium in the complex, we compared the ^31^P spectra of Li-ATP with that of a recrystallized disodium ATP trihydrate sample, prepared in a similar manner, and with existing prior data. An overlay of the ^31^P CPMAS spectra is shown in **Figure 3**. Clear shifts are observed for all signals (P_α’_ is only slightly shifted). The 2D ^31^P-^31^P DARR15 correlation experiment of recrystallized Na_2_ATP, appearing in Fig. S1 of the SI, further shows that the resonances of the γ phosphates are reversed in Li-ATP, as also can be deduced from comparison to other studies of sodium ATP trihydrate[22]. Unlike in solution, where the addition of LiCl to sodium ATP resulted in a shift of all phosphates to higher values, despite the existence of some dihydrate crystals in our data, the change in the ^31^P chemical shift is distinct, and we observe that P_α_ and P_γ’_ shift to lower chemical shift values while the rest of the phosphates shift to higher chemical shift values. The largest shift observed is for P_β_ (+1.4 ppm) followed by P_β’_ (+0.9 ppm) and P_α_ (−0.7 ppm). We will show later that these large shifts are in agreement with our observations that lithium binds to those phosphate groups. P_γ_ and P_γ’_ chemical shift changes are close to that of P_α_ (∼0.6 ppm), and they are shifted to opposite directions. The smallest change in chemical shift is observed for P_α’_.

**Figure 3.**
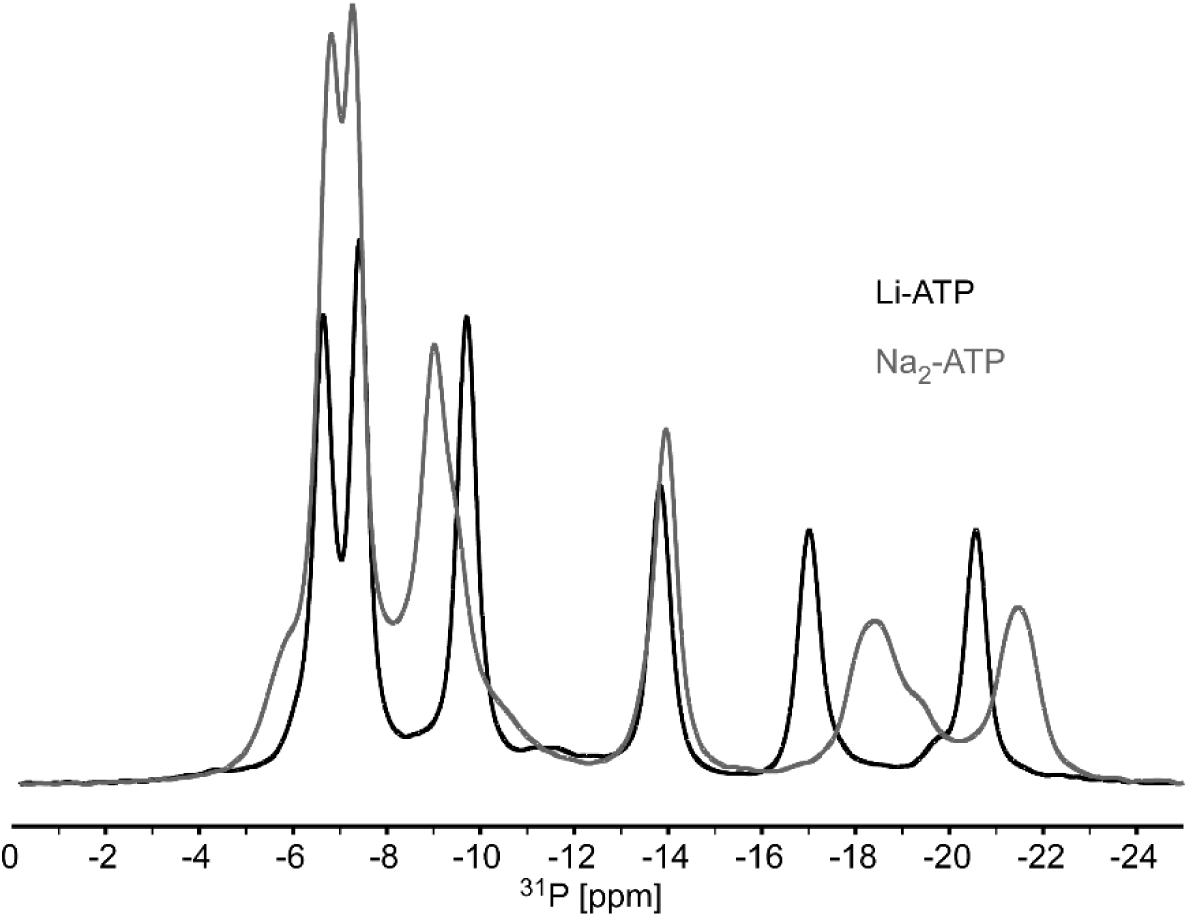
An overlay of the 1D ^31^P CPMAS spectra of Li-ATP trihydrate (black) and Na_2_ATP trihydrate (gray). Only the center bands are shown. Small contributions of the dihydrate form of Na_2_ATP can be observed as small shoulders in the spectrum of P_γ’_ and P_β_, presumably resulting from slight over-drying. Both spectra were acquired on a 9.4 T spectrometer. Li-ATP assignment from left to right: P_γ_, P_γ’_, P_α_, P_α’_, P_β_, P_β’_. The order of P_γ_ and P_γ’_ (the two peaks at ∼ -7 ppm) is reversed in Na_2_ATP.

The different trends in the chemical shift changes (both upfield and downfield) may imply on both the differences in complexation *i.e.* sodium vs. lithium, several sites vs. possibly a single metal site, changes in hydrogen bonding patterns, in the relative position of the two monomers, and on a change in the triphosphate chain conformation.

### 3.2. Direct observation of Lithium binding to ATP

Direct detection of the ^7^Li signal yielded a single (center-band) peak with a chemical shift of -0.12 ppm and a nuclear quadrupolar coupling constant of 50 kHz, determined from a SATRAS (satellite transition[42]) experiment by fitting the side band pattern of a slow spinning experiment.

In order to examine the coordination of lithium to the different phosphate groups, we performed two lithium-phosphorous heteronuclear correlation experiments. A 2D transferred echo double resonance (TEDOR[43]) experiment to observe correlations, and possibly resolve different lithium sites, and a rotational echo double resonance (REDOR[44]) experiment in order to better quantify inter-nuclear ^31^P-^7^Li distances in the complex. Both experiments have been performed following our previous optimization schemes for ^7^Li-^13^C couplings, applied to a complex of lithium, glycine and water (LiGlyW),[45] and specifically the ^31^P-^7^Li REDOR experiments follow those of van Wullen et al. performed on lithium-phosphate glasses.[37]

A ^31^P-^7^Li TEDOR spectrum is presented in **Figure 4**. ^7^Li correlations were observed with all phosphate groups save for P_α’_, which was below the noise level. Only a single lithium site is resolved, thus most probably we observe a single lithium site in the unit cell. Although the chemical shift dispersion of lithium is very small, such an experiment would have probably revealed differences between sites bound to different phosphates.

**Figure 4.**
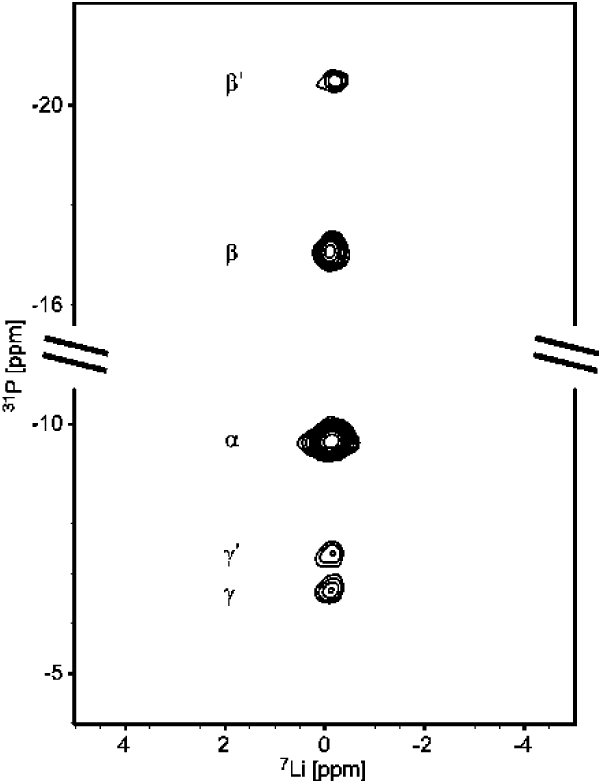
2D ^31^P-^7^Li TEDOR of {23%-^7^Li}-Li-ATP. The spectrum was acquired with a preparation mixing time of 714 μs (10 rotor periods) and a reconversion time of 1286 μs (18 rotor periods). The recoupling π pulses were applied on ^31^P at both mixing times with the exception of a single echo pulse on ^7^Li. Crosspeaks can be observed between ^7^Li at -0.15 ± 0.05 ppm and the phosphate atoms of both ATP molecules in the unit cell. No crosspeak with P_α’_ was detectable. All experimental details are given in the SI. Irradiating both ^31^P and ^7^Li, which have very close Larmor frequencies, was enabled by the use of a frequency splitter (‘REDOR-BOX’, ©NMR service GmbH).

An initial assessment of the proximity of lithium to the different ^31^P atoms can be obtained from the analysis of the cross-peak intensities, as shown in **Table 2**. Since ^31^P intensities are not similar (**Figure 1**) we normalized the TEDOR crosspeaks intensities according to the ^1^H-^31^P CP intensities in order to obtain a more accurate estimation of the transfer efficiency of each phosphate group to lithium. Lithium has the strongest correlation to P_β□_ followed by P_α_ and P_β’_, results that are in agreement with the ^7^Li-induced ^31^P shifts observed above. P_γ_ and P_γ’_ show the weakest signals, and P_α’_ is barely detected above the noise level. Lithium strongly correlating to three phosphates is in agreement with the preferred tetrahedral coordination for lithium, as we observed in LiGlyW[45] (correlations to carboxyl groups), assuming that the fourth ligand is a water molecule, a base nitrogen, or from the ribose.

**Table 2.** ^31^P CP relative intensities and ^31^P-^7^Li TEDOR normalized crosspeak intensities in Li-ATP. All intensities were normalized according to P_γ’_.

| | $\text{P}_{\alpha}$ | $\text{P}_{\beta}$ | $\text{P}_{\gamma}$ | $\text{P}_{\alpha'}$ | $\text{P}_{\beta'}$ | $\text{P}_{\gamma'}$ |
| --- | --- | --- | --- | --- | --- | --- |
| $^1\text{H}$-$^{31}\text{P}$ CP | 1.10 | 2.64 | 1.19 | 2.66 | 2.69 | 1.00 |
| TEDOR | 3.85 | 5.05 | 1.45 | - | 3.15 | 1.00 |

In order to better quantify the coordination of lithium, we performed CPMAS-filtered ^31^P{^7^Li} REDOR experiments in order to extract ^7^Li-^31^P distances. All REDOR curves are presented in **Figure 5**. It can be seen that the three phosphate groups P_β_, P_α_ and P_β’_ show the strongest recoupling, and have a similar initial slope indicating a comparable inter-nuclear distance. The other three phosphate groups also show a comparable recoupling curve however they are clearly more distant from the lithium ion.

**Figure 5.**
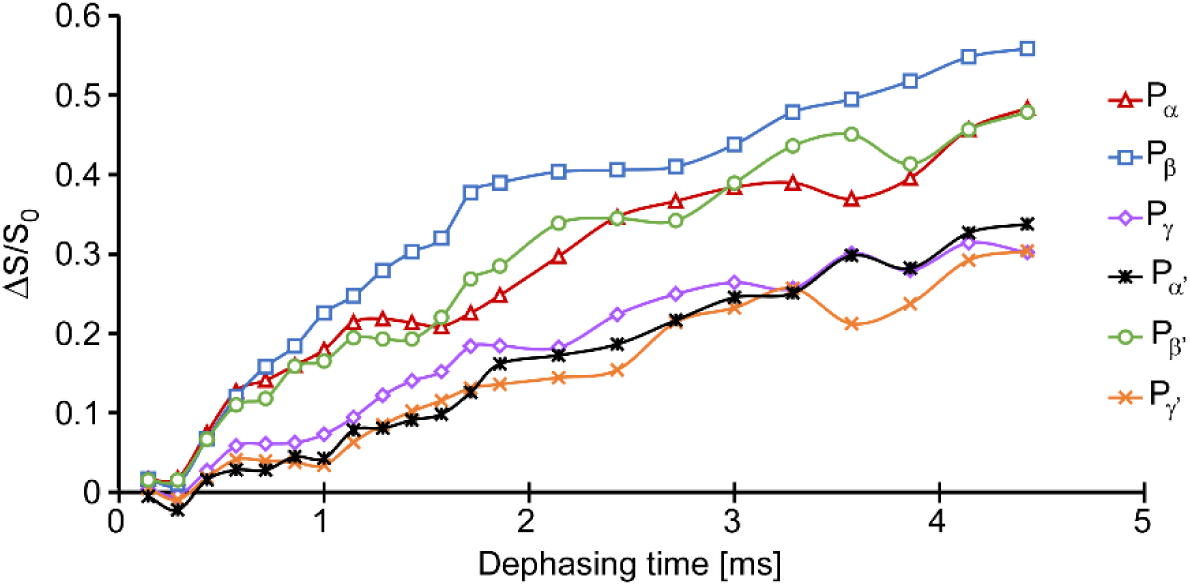
Experimental ^31^P-detected CP-filtered ^31^P{^7^Li} REDOR dipolar recoupling curves for the six non-equivalent ^31^P sites in {23%-^7^Li}-Li-ATP. In the REDOR sequence all dephasing pulses were applied to the ^7^Li channel due to the relatively large CSA of the ^31^P spins. P_α_, P_β_ and P_β’_ are closest to the lithium center. The REDOR fraction ΔS/S_0_ is given by (*S*_*0*_-*S*)/*S*_*0*_ where *S*_*0*_ is a reference signal obtained from the intensity of the ^31^P spectrum obtained without the pulses on lithium, and *S* is the signal with all pulses.

In order to quantify the inter-nuclear ^31^P-^7^Li distances we first examined the effect of various multi-spin systems on the signals, and assumed a single lithium site, since we can only resolve a single lithium signal, even when attempting to separate the sites with a triple-quantum/single quantum ^7^Li correlation experiment, as done before for lithium in the enzyme Inositol monophosphatase[46]. Simulations of a LiP_*n*_ spin system with *n*=1-3 were performed taking into account the nuclear quadrupolar coupling constant Cq of lithium, and CSA values of the phosphates, obtained from our experimental data. ^31^P-^31^P homonuclear dipolar couplings, and Euler angles relating the CSA and dipolar tensors were extracted from the structure of Na_2_ATP[22]. The simulations (SI, **Figure S2**) showed that the initial rise (up to about 65% of the total possible recoupling value) of the REDOR curve (from which the inter-nuclear distance can be measured) was unaffected by the spin system size or by the Euler angles of the interaction tensors. Small oscillations were observed at longer dephasing times and do not affect the analysis of the initial rise. Similar observations have been made before by Bertmer and Eckert,[47] who used the identity of the initial rise to fit multi-spin systems using second moment analysis. Thus we concluded that considering a two spin system was sufficient to simulate the initial rise of the REDOR dipolar recoupling curve.

The experimental data for the six ^31^P species shown in **Figure 5** were then deconvoluted (using the software DMFIT[41]) and fit individually to simulations at a range of ^31^P-^7^Li distances using a single spin pair. **Figure 6** shows on top the results for the closest ^31^P spins, P_α_, P_β_, and P_β’_. All curves are in agreement with a distance of 2.9-3.1 Å (with errors of 0.3-0.4 Å). The bottom shows the results for P_α’_, P_γ_, and P_γ’_, and they report on distances of 3.4-4.1 Å with a similar error range. Reported metal-phosphorus distances in ATP complexes are 3.4-3.8 Å in sodium ATP[23–25] and 3.1-3.3 Å in magnesium ATP[30,31]. Thus, the distances in Li-ATP are in a better agreement with the MgATP complex. The current results of oxygen-mediated ^7^Li-^31^P are also in agreement with prior observations of oxygen-mediated ^7^Li-^31^C distances including our studies of the LiGlyW complex,[45,48] and of lithium binding in the enzyme inositol monophosphatase.[46] Thus they represent typical Li-O-X coordination.

**Figure 6.**
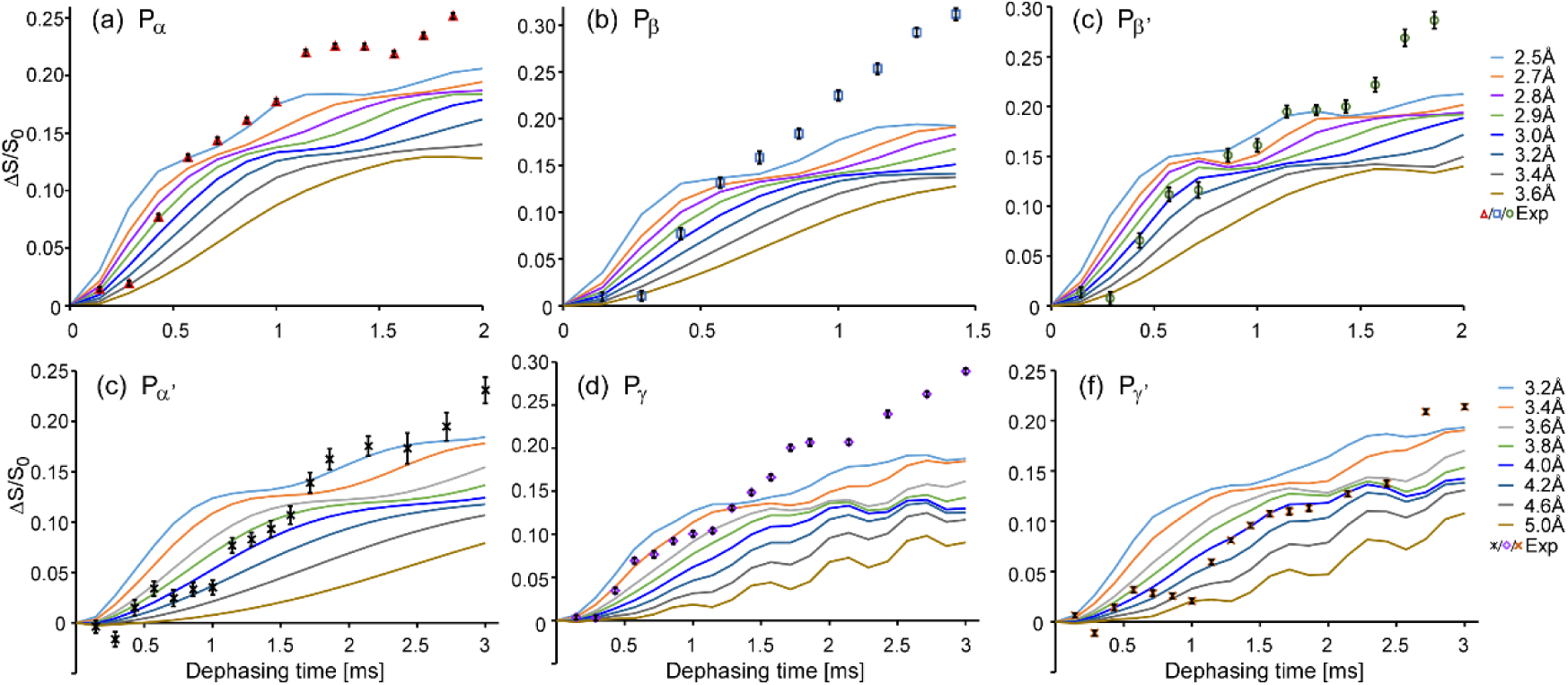
Comparison of experimental CP-filtered, ^31^P-detected REDOR data fractions (δ*S*/*S*_*0*_) of the close phosphate signals (a) P_α_, (b) P_β_, (c) P_β’_, and the more remote phosphates (d) P_α’_, (e) P_γ_ and (f) P_γ’_ with simulations of single Li−P spin pairs. The simulations were multiplied by 0.23 in order to account for the labeling pattern and the natural ^7^Li percentage. The sample is {23%-^7^Li}-Li-ATP. The best fits for the close phosphates, extracted from the initial rise (up to ∼0.6 ms) are for P_α_ 2.9±0.35Å (D=780 Hz), for P_β_ 3.0±0.45Å (D=700 Hz), and for P_β’_ 3.1±0.3Å (D=630 Hz). For the remote phosphates (d-f, fit up to ∼1.5 ms), the best fit for P_α’_ is 4.0±0.35Å (D = 300 Hz), for P_γ_ is 3.4±0.3Å (D = 480 Hz) and for P_γ’_ is 4.1±0.4Å (D = 270 Hz). The best fit distances were extracted from the RMSD plots shown in Figure S3 of the SI. The deviation of the experimental points beyond 1-1.5 ms occurs due to the existence of remote ^7^Li labeled sites. The oscillatory nature of the curves for P_γ/γ’_ is probably caused by off resonance effects (P_α’_ is on resonance and P_γ/γ’_ has an offset of 6-7 ppm).

While the data above are sufficiently reliable to provide clear evidence on the proximity of lithium to the various phosphate moieties, several errors may marginally affect the quantification of our data. As discussed above, the initial rise is the most important region for a successful fit, however, the number of data points in this region for the shorter distances is small due to (i) the hardware limitation of using of a moderate spinning speed of 14 kHz (hence longer times between the detection points); (ii) experimental deviations for data points taken when a full XY4 or XY8 refocusing cycle could not be used, evident for the second data point in Figure 6. Yet, these do not affect the conclusions -

From ^31^P and ^7^Li shifts, from ^31^P-^7^Li correlation NMR experiments, and from ^31^P-^7^Li distance measurement experiments, clearly a single lithium site is coordinated to P_α_, P_β_ and P_β’_, with inter-nuclear ^31^P-^7^Li distances of ∼ 3Å.

### 3.3. ^1^H NMR and the spatial organization of Li-ATP

In order to further characterize the triphosphate chain and the coordination of lithium in the Li-ATP complex, we performed 2D heteronuclear correlation experiments involving protons. Both ^1^H–^31^P *w*PMLG–HETCOR spectra and ^1^H-detected {^1^H}^31^P-^1^H spectra performed at a MAS frequency of 62 kHz, shown in **Figure 7**, show signals of acidic protons and of other protons belonging to the adenine base and the ribose. Identification of the signals is in part supported by the solution NMR spectrum shown and assigned in the SI (**Figure S4**). Two strong signals, resonating at 11.6 ppm and 15.3 ppm are missing from the solution spectrum, and exist in the MAS spectrum. A third, weaker signal appears only in the MAS spectrum acquired at 62 kHz. Those signals belong to exchangeable acidic protons. Their assignment is based on their relative intensities at different mixing times; the signal resonating at 11.6 ppm shows strong correlations to the gamma phosphates, and has very fast build-up rates (see SI, **Figure S5**). It also shows correlations to P_α_ and P_α’_ although with a lower intensity. This signal therefore corresponds to the acidic P_γ_ phosphate groups. Indeed such shifts are typical of hydrogen bonded protonated phosphates[49]. The correlations to the alpha phosphates are indicative of the spatial organization of the tri-phosphate moiety locating P_γ_ of one ATP molecule between P_α’_ and P_γ’_ of a second ATP molecule. In our preparation of the Li-ATP, the pH value was approximately 2-3, and following previous studies[50–53] indicating that one hydroxyl group of the gamma phosphate has a pKa of ∼1-2 and the second group has a pKa of ∼6-7, we conclude that only one of the gamma phosphate hydroxyl groups is protonated. Thus, the other low-field signal that shows strong correlations to primarily P_γ_ and P_γ’_ at 15.3 ppm, albeit with a smaller intensity, does not belong to a phosphate group. Unlike the signal at 11.6 ppm, the correlation to P_α_ is much weaker and correlations to both P_β/β’_ phosphates appear as well.

**Figure 7.**
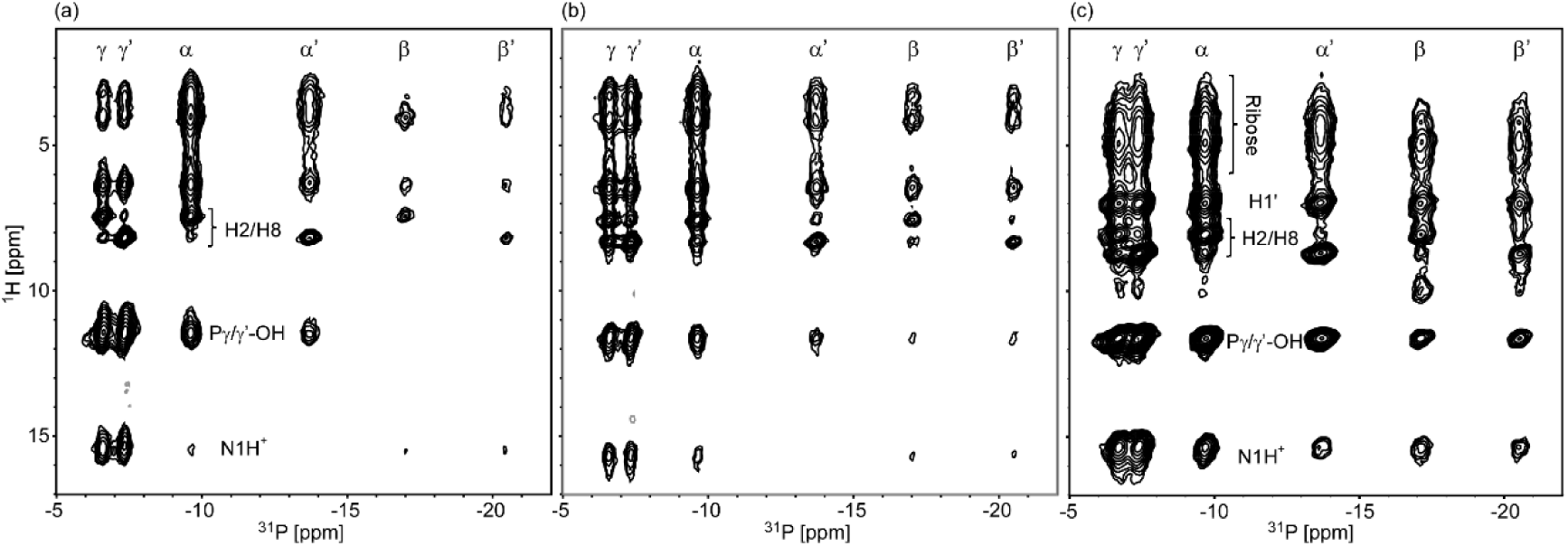
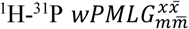 [56,57] HETCOR spectra of Li-ATP acquired at a spinning speed of 11.5 kHz (at 0°C) and CP contact times of (a) 1 ms and (b) 3 ms. The ^1^H axis is scaled. (c) ^1^H-detected {^1^H}^31^P-^1^H HETCOR MAS spectrum of Li-ATP acquired at a spinning speed of 62 kHz (at -30°C) with a ^31^P→^1^H CP contact time of 2.75 ms. The spectrum was flipped in order to align the axes of all spectra in the Figure. The final ^1^H chemical shifts were determined according to the ^1^H-detected experiment (c). A full list of experimental parameters appears in the SI. Most probable ^1^H assignment are indicated on plot (c) and tabulated in the SI.

Since the pKa of N1 is ∼4-5[50,51], it is protonated in our preparation. Imino NH protons involved in Watson-Crick hydrogen bonding resonate at 12-14 ppm, and in isocytosine and guanosine-derivatives for example,[54,55] they have chemical shifts of 13-15 ppm. Thus the peak at 15.3 ppm can be attributed to the imino proton attached to N1 and is probably subjected to strong hydrogen bonding. The strong correlations to P_γ/γ’_ suggest a bending of the base towards the edge of the triphosphate of the second ATP molecule. Approximate similar intensities of crosspeaks to P_α/α’_, and to P_β/β’_ correspond to the location of the ring mid-way between its own P_α_ (or P_α’_) and the P_β’_ (or P_β_) of the second ATP molecule.

Additional proton correlation experiments have been performed with lithium. A *w*PMLG-HETCOR experiment acquired with a short mixing time (**Figure 8a**) does not reveal contacts of lithium with the acidic protons, further strengthening our conclusion that lithium is not in proximity to P_γ_. Such correlations only appear at a longer mixing time (**Figure 8b,** ^1^H-detected {^1^H}^7^Li-^1^H HETCOR at 62 kHz MAS). The projection of the ^1^H spectrum reveals a broad hump underneath the spectrum, which was not apparent in the ^31^P-^1^H spectrum. Most probably this signal corresponds to a water molecule (or several) that coordinates lithium, completing its fourth coordination. Correlations with some of the ribose signals appear at 4-7 ppm, which cannot be faithfully assigned at this stage, however are in line with the fact that lithium coordinates P_α_ and is thus in proximity to those hydrogens.

**Figure 8.**
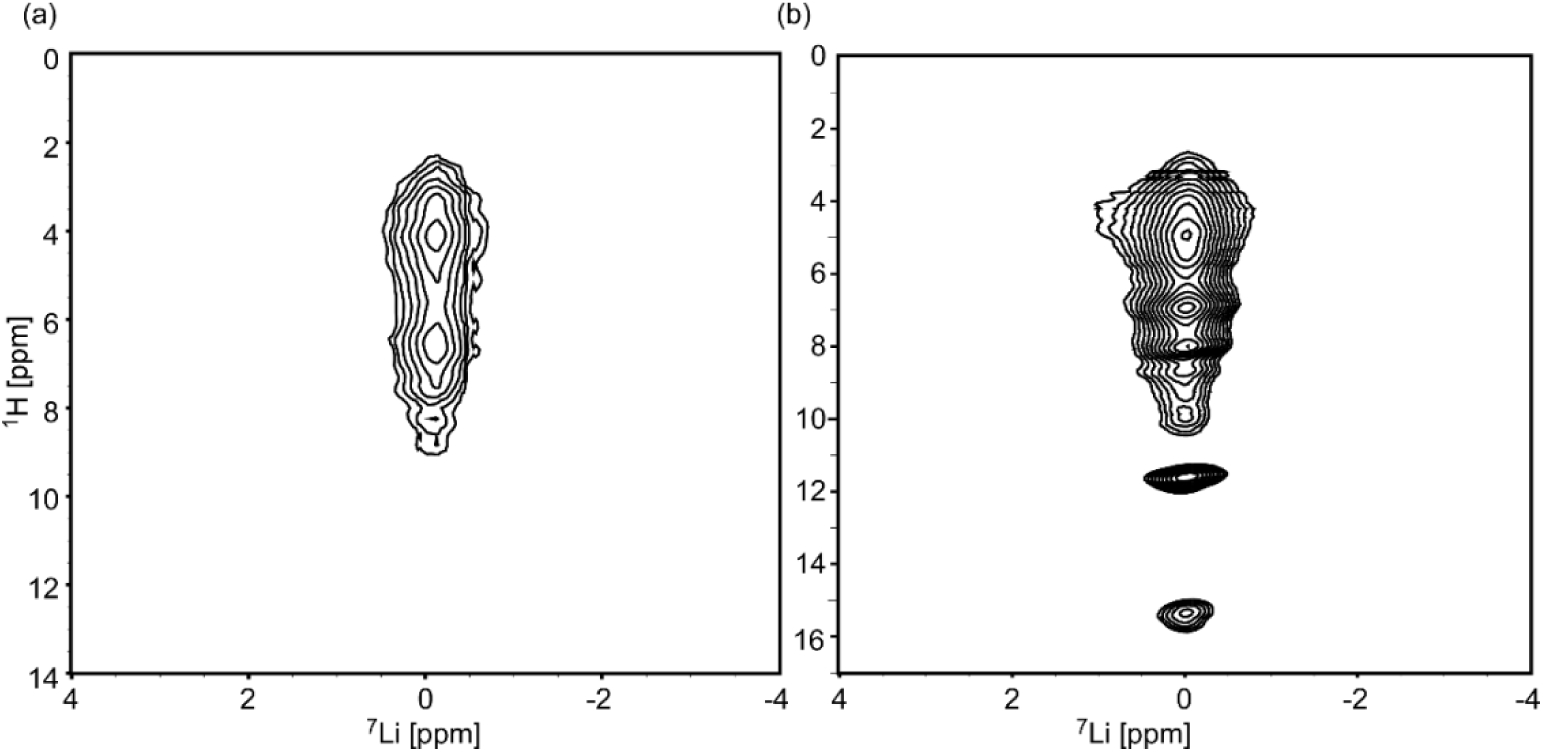
(a) 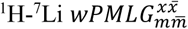 HETCOR spectrum of {23%-^7^Li}-Li-ATP acquired at a spinning speed of 11.5 kHz (at -20°C), and a CP contact time of 0.5 ms. The ^1^H axis is scaled. (b) {^1^H}^7^Li-^1^H HETCOR of {NA-^7^Li}-Li-ATP acquired with a CP contact time of 1.7 ms (62 kHz at -30°C). The spectrum was flipped in order to align the axes of all spectra in the Figure. The final ^1^H chemical shifts were determined according to the ^1^H-detected experiment at 62 kHz. A full list of experimental parameters appears in the SI.

Potential assignments of the ^1^H signals, based on ^1^H-X correlations, on ^1^H-^31^P build-up curves, and on the ^1^H solution NMR spectrum of Li-ATP appear in the SI.

### 3.4. Residual sodium in Li-ATP

The Li-ATP complex was prepared from mixing excess LiCl with commercial disodium ATP trihydrate, in the molar ratio Li:Na of 10:1. Therefore, sodium may remain as a complex of the form NaLi-ATP, or as a residual with no specific role in the complex. In the trihydrate form of Na_2_ATP there are four chemically inequivalent sites according to X-ray crystallography[24], as well as multiple-quantum MAS NMR[29] (also observed by us, see SI, **Figure S6**). An overlay of 1D MAS ^23^Na spectra of Li-ATP and of recrystallized Na_2_ATP are shown in **Figure 9**. Clearly some of the sodium signal disappears in the 1D ^23^Na spectrum of Li-ATP, however, two signals (at least) remain; a narrow peak and a broadened signal. MQMAS spectra revealed three sites (**Figure S6**) and a three-site-fit for the 1D experiment generated Cq values of approximately 0.85, 2.3 MHz and 0.95 MHz, respectively, where the third site has a low occupancy in comparison to the other two.

**Figure 9.**
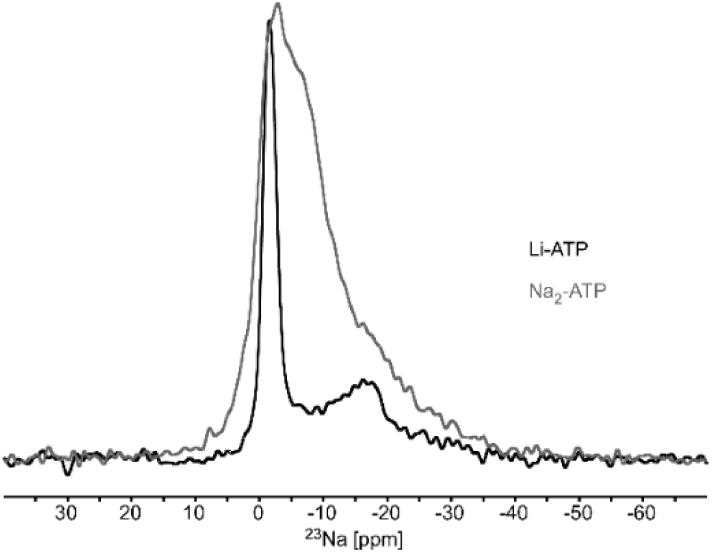
Overly of 1D ^23^Na spectra of Li-ATP (black) and Na_2_ATP (gray). MQMAS spectra of both samples appear in the SI, Fig. S6.

In order to further characterize these sodium sites, we performed ^7^Li{^23^Na} and ^31^P{^23^Na} PM-RESPDOR[58,59] distance measurement experiments. A ^7^Li{^23^Na} PM-RESPDOR recoupling curve is shown in **Figure 10**. We observe a clear recoupling of the lithium signal that is consistent with an inter-nuclear distance of 4.3Å, and the recoupling approaches the theoretical limit for a spin-3/2 of ΔS/S_0_=0.75. Therefore, it is most likely that at least one site in every ATP molecule can be attributed to sodium in close proximity to lithium. Since MQMAS suggests that at least two sodium sites exist in every ATP dimer, we can assume that both are in similar proximity to lithium and a simulation with two sodium sites shows even a better fit (at longer times) than a single spin-pair, and the estimated distance then increases up to 5Å. In both cases, every ATP molecule must have a persistent high affinity sodium site.

**Figure 10.**
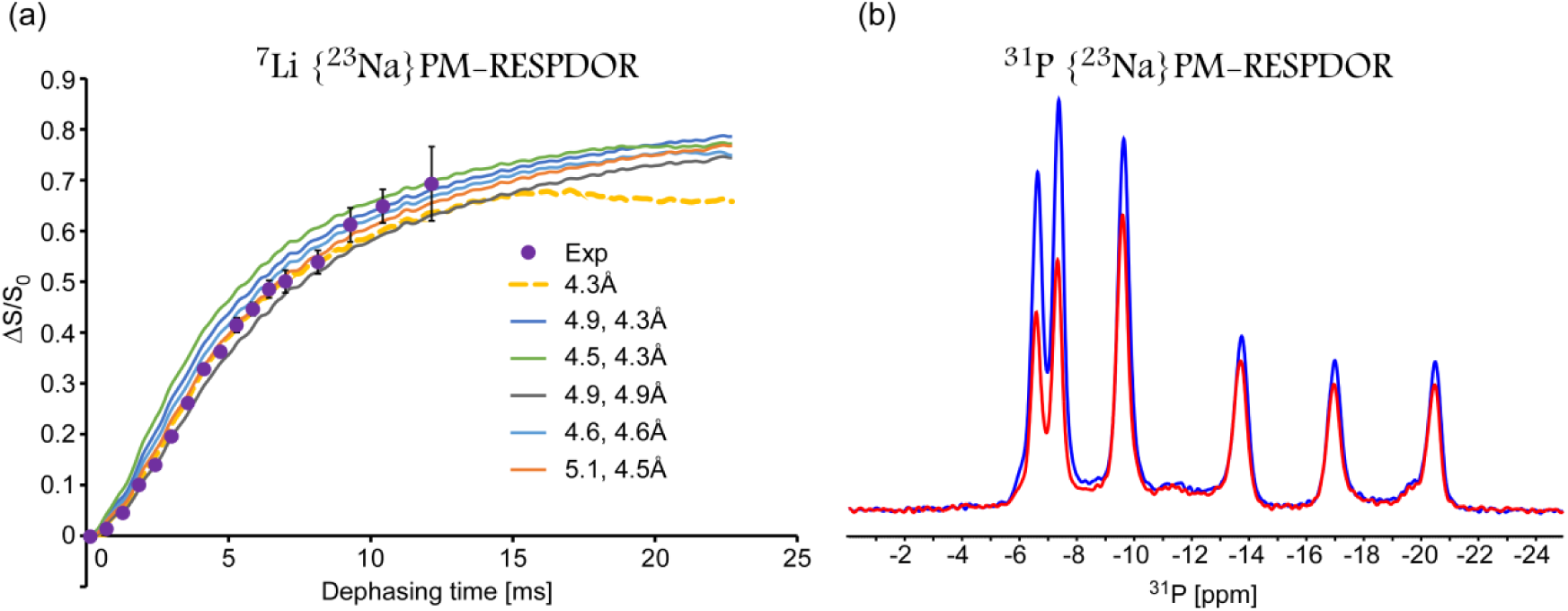
(a) PM-RESPDOR ^7^Li{^23^Na} recoupling curve of Li-ATP. A fit to an isolated pair yields a lithium-sodium distance of 4.3Å (dash line, D = 150 Hz). The experimental points ΔS/S_0_ are shown as circles. SIMPSON simulated curves for two sodium sites at various distances up to 5.1Å are shown and the experimental data are in good agreement with D = 90, 130 Hz (orange line, 4.5Å, 5.1Å). (b) A single ΔS/S_0_ point on the PM-RESPDOR ^31^P{^23^Na} recoupling curve, acquired at a mixing time of 1.3 ms, showing the S_0_ reference signal (blue) and the decay signal S (red). From left to right in the spectrum of Li-ATP, P_γ_ and P_γ’_ have the most pronounced initial decay, followed by P_α_. A much slower decay is observed for the other phosphates (see also Figure S7 in the SI for the complete curves).

Furthermore, looking at one point in a ^31^P{^23^Na} PM-RESPDOR experiment (**Figure 10b**) reveals that the sodium atom coordinates to P_γ_, P_γ’_, and to a lower extent, to P_α_. The complete recoupling curves are shown in the SI, **Figure S7**. Assuming a decay from a single sodium spin, fitting the ^31^P signal of P_γ_, is consistent with a distance of 3Å, which is too short when comparing to the reported Na-P distances in Na_2_ATP.[23–25] A fit involving two sodium sites yields more plausible distances of 3.3-3.4 Å.

Another way to directly determine if the additional sodium sites are all part of the structure, and to characterize their environment, is to perform sodium-detected REDOR experiments. We therefore implemented ^23^Na{^31^P} REDOR and ^23^Na{^7^Li} REDOR experiments. We observe that in both experiments (**Figure 11** and **Figure S8**) all sodium sites decay with an approximately equal rate. Thus, there must be at least two inequivalent sodium sites in the sample (and a possible third site). The ^23^Na{^7^Li} REDOR experimental results in **Figure 11** show that sodium recoupling reaches ∼0.5 (with 24% dilution of ^7^Li) rather than the theoretical 0.24, suggesting that every sodium atom is in contact with two lithium sites. A two spin system simulation shows a reasonable fit to a distance of 5.5±0.5Å. This is in agreement with the inverse ^7^Li{^23^Na} PM-RESPDOR measurements.

**Figure 11.**
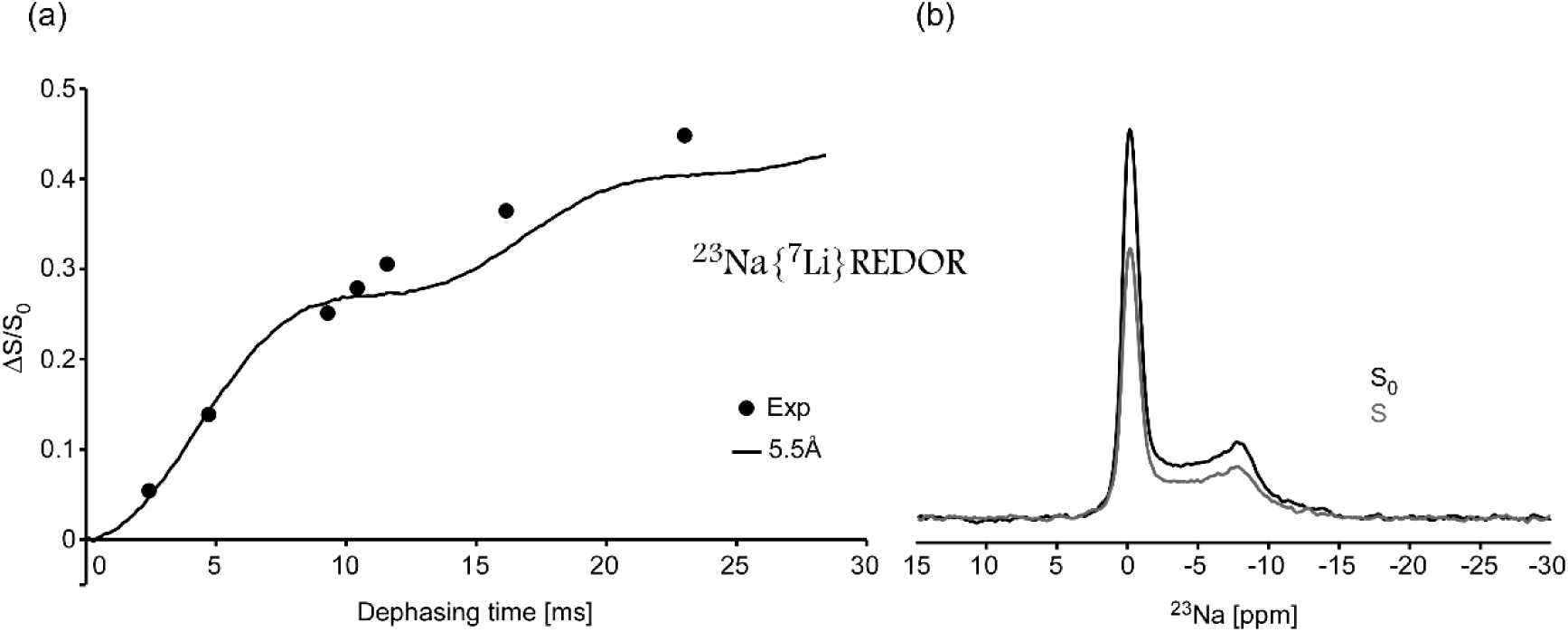
(a) ^23^Na{^7^Li} REDOR recoupling curve in Li-ATP. Data points represent the integral of the entire sodium signal. A SIMPSON simulated curve for a Li-Na distance of 5.5Å (D = 70 Hz) is shown. (b) A single ΔS/S_0_ point acquired at a mixing times of 11.57 ms showing the S_0_ reference signal (black) and the decay signal S (gray). It can be seen that the dephasing seems to be uniform.

### 3.5. Proposed schematic model for lithium binding to ATP

Based on the results of this work we are able to propose the schematic model for lithium binding in the lithium ATP complex shown in **Figure 12**. In the solid phase, Li-ATP is a dimer, similarly to Na_2_ATP (MgATP is a dimer of two symmetrically equivalent molecules). The arrangement is such that P_γ_ of one molecule is located between P_α’_ and P_γ’_ of a second ATP molecule, closer to P_α’_, and with P_β’_ located at an angle to both. This bent arrangement also occurs in the other structures, shown in Figure S9, and is a result of the restricted P-O-P bond angle. Yet, the relative arrangement of the two molecules must be different from both Na_2_ATP and MgATP/BPA as revealed by our ^31^P-^31^P correlation spectra (see also Section 3.1).

**Figure 12.**
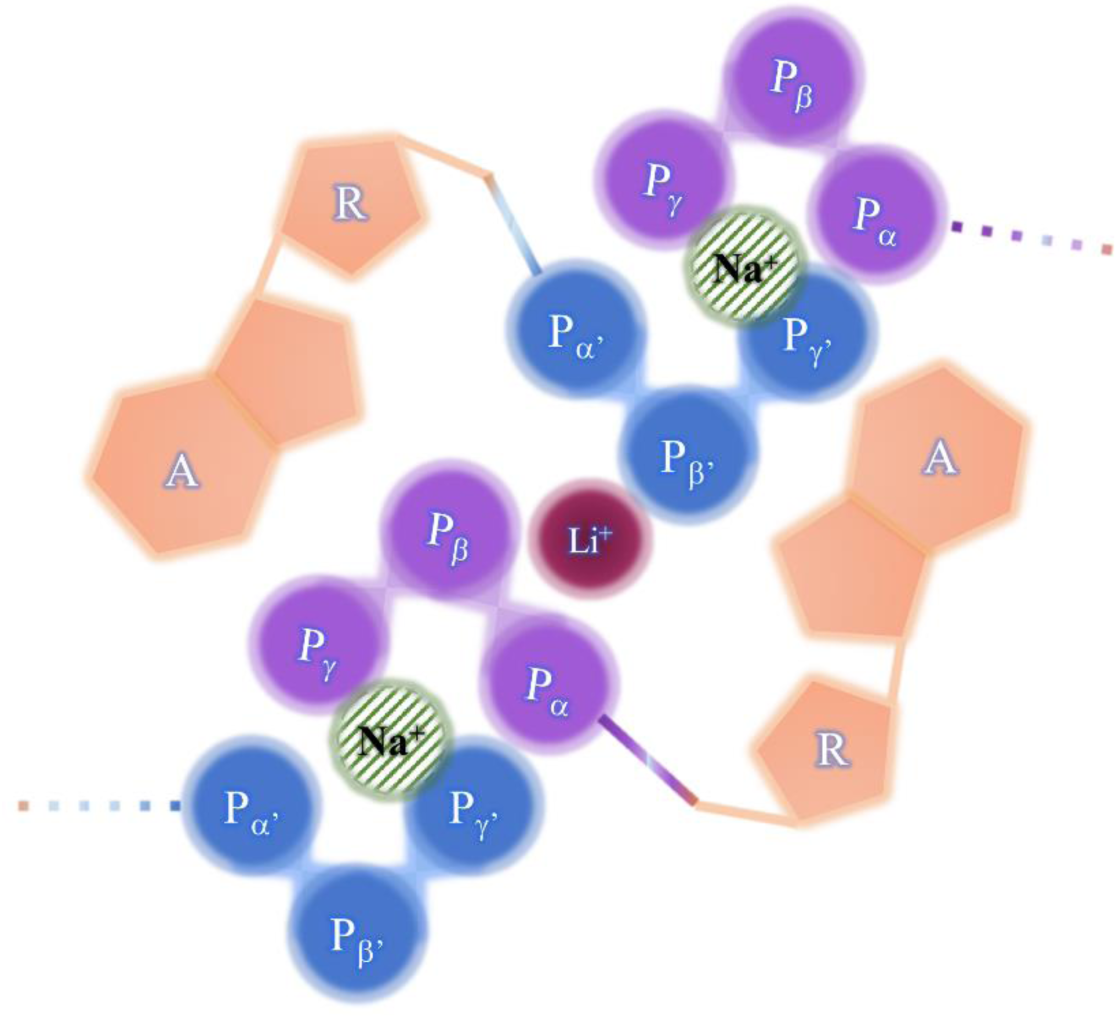
A schematic model for lithium binding to two ATP molecules via three phosphate groups.

Lithium is coordinated to the three phosphates P_α_, P_β_ and P_β’_ at a distance of 3.0(±0.4)Å, and a forth ligand is presumably a water molecule. The adenine ring is somewhat tilted positioning its imino proton N_1_H^+^ towards the gamma phosphate of a second ATP molecule.

Every such complex also has at least two sodium cations despite the fact that lithium was used in excess. These sodium ions are located at a distance of 4.3 – 5.5 Å from two lithium sites (both at equivalent crystallographic positions), and are coordinated to the phosphates P_γ_ and P_γ’_, with P_α_ the next closest phosphate. Such binding is observed for Na1 and Na2 in the structure of Na_2_ATP, and hence may be those that are preserved in Li-ATP. No analogue can be found in MgATP, where one Mg^2+^ site coordinates six water molecules, and the other three phosphates from each ATP molecules. Yet, Li-P distances are similar to Mg-P (3.15-3.28Å), and not to Na-P (3.5-3.7 Å).

Despite the fact that the model is not complete in the sense that we cannot determine atomic coordinates at this stage, it demonstrates well the coordination of lithium, the strong affinity and position of the sodium sites, and the overall organization of the two ATP molecules.

A more elaborate atomic-resolution model requires additional experiments and probably a full NMR crystallography approach.

## 4. Summary and Conclusions

Lithium in the form of a salt (e.g. Li_2_CO_3_) is one of the most prominent drugs for various mental illnesses, mainly bipolar disorder. Lithium binds many enzymes as well as ATP, “the chemical energy of life”. In this paper we characterize the complex of lithium with ATP in a crystalline solid using a multi-nuclear MAS solid-state NMR approach. The experiments included simple one-dimensional spectra, two-dimensional correlation experiments involving lithium, phosphates and sodium, ^1^H correlation experiments, and explicit distance measurements.

Based on all the accumulated data we can define the lithium coordination and identify the existence of structural sodium ions, bound to the complex in high affinity. To the best of our knowledge, an experimental model based on direct observation of lithium does not exist, while other models in solution suggest binding to gamma phosphates. Here we show a model of binding for the Li-ATP complex, in which lithium coordinates to the three phosphate groups P_α_, P_β_ and P_β’_ with equivalent distances, and to a water molecule. Li-P distances are similar to those in MgATP/BPA[18], to Li-C in a lithium-glycine-water complex[45], and similar to Li-C in the enzyme inositol monophosphates, the putative target of lithium therapy[46]. This is a result that stems from the fact that Li^+^ and Mg^2+^ have a similar ionic radii.[60] The methods we present here are suitable for characterization of metal-bound ATP molecules, even when a mixture of cations exist, a state depicting more biologically relevant situations.

While our model depicts lithium coordination in a crystalline environment, it may well be related to ATP-bound enzymes. For example, Sugawara et al.[25] have shown that one of the ATP molecules in the unit cell of Na_2_ATP resembles that in enzymes. Moreover, since lithium binds two phosphates of one ATP molecules, and a third from a second molecule, in a physiological environment the second molecule may be replaced with a carboxyl group (considering that C-O-Li distances are similar P-O-Li distances[45]), or a similar moiety. In addition, the high hydration of solid metal-ATP complexes is also consistent with such environments in the cell.

## 5. Author contributions

A.G and A.H.: Conceptualization, methodology, investigation, writing, visualization; A.G.: Funding acquisition.

## 6. Acknowledgement

This work was supported by the US-Israel Binational Science Foundation grant #2016169.

## References

[1] W. Young, Review of Lithium Effects on Brain and Blood, Cell Transplant., 18 (2009) 951–975.

[2] H. Eldar-Finkelman, Glycogen synthase kinase 3: an emerging therapeutic target, Trends Mol. Med., 8 (2002) 126–132.

[3] J.A. Quiroz, T.D. Gould, H.K. Manji, Molecular effects of lithium, Mol. Interv., 4 (2004) 259–272.

[4] E. Jakobsson, O. Argüello-Miranda, S.-W. Chiu, Z. Fazal, J. Kruczek, S. Nunez-Corrales, S. Pandit, L. Pritchet, Towards a Unified Understanding of Lithium Action in Basic Biology and its Significance for Applied Biology, J. Membr. Biol., 250 (2017) 587–604.

[5] R.S. El-Mallakh, Ion homeostasis and the mechanism of action of lithium, Clin. Neurosci. Res., 4 (2004) 227–231.

[6] K. Burton, Formation Constants for the Complexes of Adenosine Di- or Tri-phosphate with Magnesium or Calcium Ions, Biochem. J., 71 (1959) 388–395.

[7] L. Nanninga, The association constant of the complexes of adenosine triphosphate with magnesium, calcium, strontium, and barium ions, Biochim. Biophys. Acta, 16 (1961) 330–338.

[8] W.J. O’Sullivan, D.D. Perrin, The Stability Constants of Metal-Adenine Nucleotide Complexes, Biochemistry, 3 (1964) 18–26.

[9] M.S. Mohan, G.A. Rechnitz, Ion-Electrode Study of Magnesium(II)-ATP and Manganese(II)-ATP Association, Arch. Biochem. Biophys., 162 (1974) 194–199.

[10] R. Adolfsen, E.N. Moudrianakis, Control of Complex Metal Ion Equilibria in Biochemical Reaction Systems, J. Biol. Chem., 253 (1978).

[11] J. Wilson, A. Chin, Chelation of divalent cations by ATP, studied by titration calorimetry, Anal. Biochem., 19 (1991) 16–19.

[12] M.M. Taqui Khan, A.E. Martell, Metal Chelates of Adenosine Triphosphate, 66 (1962) 10–15.

[13] E.O. Bishop, S.J. Kimber, D. Orchard, B.E. Smith, A 31P-NMR study of mono- and dimagnesium complexes of adenosine 5′-triphosphate and model systems, BBA - Bioenerg., 635 (1981) 63–72.

[14] R. Prigodich, P. Haake, Association phenomena 5 Association of cations with nucleoside di-and triphosphates studied by phosphorus-31 NMR, Inorg. Chem., (1985) 89–93.

[15] A. Abraha, D.E.M. Defreitas, M. Margarida, C.A. Castro, C. Geraldes, Competition between Li+ and Mg2+ for ATP and ADP in aqueous-solution - a multinuclear NMR-study, J. Inorg. Biochem., 42 (1991) 191–198.

[16] M. Cohn, T.R. Hughes, Phosphorus Magnetic Resonance Spectra of Adenosine Di- and Triphosphate II Effect of Complexing with Divalent Metal Ions, J. Biol. Chem., 237 (1962) 176–182.

[17] T.-D. Son, M. Roux, M. Ellenberger, Interaction of Mg2+ ions with nucleoside triphosphates by phosphorus magnetic resonance spectroscopy, Nucleic Acids Res., 2 (1975) 1101–1110.

[18] F. Ramirez, J.F. Marecek, Coordination of Magnesium with Adenosine 5’-Diphosphate and Triphosphate, Biochim. Biophys. Acta, 589 (1980) 21–29.

[19] J.L. Bock, The binding of metal ions to ATP: a proton and phosphorus nmr investigation of diamagnetic metal--ATP complexes, J. Inorg. Biochem., 12 (1980) 119–30.

[20] L. Amari, B. Layden, Q.F. Rong, C. Geraldes, D.M. de Freitas, Comparison of fluorescence, P-31 NMR, and Li-7 NMR spectroscopic methods for investigating Li+/Mg2+ competition for biomolecules, Anal. Biochem., 272 (1999) 1–7.

[21] Y. Shindo, A. Naito, S. Tuzi, Y. Sugawara, Stepwise conformational transition of crystalline disodium adenosine 5′-triphosphate with relative humidity as studied by high resolution solid state 13, J. Mol. Struct., 603 (2002) 389–397.

[22] M.J. Potrzebowski, J. Gajda, W. Ciesielski, I.M. Montesinos, Distance measurements in disodium ATP hydrates by means of P-31 double quantum two-dimensional solid-state NMR spectroscopy, J. Magn. Reson., 179 (2006) 173–181.

[23] O. Kennard, N.W. Isaacs, W.D.S. Motherwell, J.C. Coppola, D.L. Wampler, A.C. Larson, D.G. Watson, The Crystal and Molecular Structure of Adenosine Triphosphate, Proc. R. Soc. A Math. Phys. Eng. Sci., 325 (1971) 401–436.

[24] a. C. Larson, Restrained refinement of disodium adenosine 5’-triphosphate trihydrate, Acta Crystallogr. Sect. B Struct. Crystallogr. Cryst. Chem., 34 (1978) 3601–3604.

[25] Y. Sugawara, N. Kamiya, H. Iwasaki, T. Ito, Y. Satow, Humidity-Controlled Reversible Structure Transition of Disodium Adenosine 5’-Triphosphate between Dihydrate and Trihydrate in a Single Crystal State, J. Am. Chem. Soc., (1991) 5440–5445.

[26] S. Ding, C. McDowell, 23Na Solid-state NMR studies of hydrated disodium adenosine triphosphate, Chem. Phys. Lett., (2000) 316–322.

[27] A. Wong, G. Wu, Characterization of the Pentacoordinate Sodium Cations in Hydrated Nucleoside 5’-Phosphates by Solid-State 23Na NMR and Quantum Mechanical Calculations, J. Phys. Chem. A, (2003) 579–586.

[28] C. V Grant, D. McElheny, V. Frydman, L. Frydman, Solid-state NMR investigation of sodium nucleotide complexes, Magn. Reson. Chem., 44 (2006) 366–74.

[29] M. Nausner, J.J. Brus, M. Haeubl, N. Mueller, W. Schoefberger, M. Häubl, N. Müller, Characterization of the sodium binding sites in microcrystalline ATP by Na-23-solid-state NMR and ab initio calculations, Inorganica Chim. Acta, 362 (2009) 1071–1077.

[30] R. Cini, M.C. Burla, A. Nunzi, G.P. Polidori, P.F. Zanazzi, Preparation and physico-chemical properties of the ternary complexes formed between adenosine 5′-triphosphoric acid, bis(2-pyridyl)amine, and divalent metal ions Crystal and molecular structures of the compounds containing Mg II and Ca II, J. Chem. Soc., Dalt. Trans., (1984) 2467–2476.

[31] G. Tamasi, F. Berrettini, M.B. Hursthouse, R. Cini, Effect of Free Water Molecules on the Structure of Mg-ATP-Dipyridylamine and Overview on Selected Metal-Adenosine Triphosphate Structures in Model Compounds and in Enzymes, Open Crystallogr. J., 3 (2010) 1–13.

[32] R. Cini, M. Sabat, M. Sundaralingam, M.C. Burla, A. Nunzi, G. Polidori, P.F. Zanazzi, Interaction of Adenosine 5′-Triphosphate with Metal Ions X-ray Structure of Ternary Complexes Containing Mg(II), Ca(II), Mn(II), Co(II), ATP and 2,2′ -Dipyridylamine, J. Biomol. Struct. Dyn., 1 (1983) 633–637.

[33] C. V. Grant, V. Frydman, L. Frydman, Solid-State 25 Mg NMR of a Magnesium(II) Adensosine 5‘-Triphosphate Complex, J. Am. Chem. Soc., 122 (2000) 11743–11744.

[34] D. Mota de Freitas, M.M.C.A. Castro, C.F.G.C. Geraldes, Is Competition between Li + and Mg 2+ the Underlying Theme in the Proposed Mechanisms for the Pharmacological Action of Lithium Salts in Bipolar Disorder?, Acc. Chem. Res., 39 (2006) 283–291.

[35] K.T. Briggs, G.G. Giulian, G. Li, J.P.Y. Kao, J.P. Marino, A Molecular Model for Lithium’s Bioactive Form, Biophys. J., 111 (2016) 294–300.

[36] T. Dudev, C. Grauffel, C. Lim, How native and alien metal cations bind ATP: Implications for lithium as a therapeutic agent, Sci. Rep., 7 (2017) 1–10.

[37] L. van Wüllen, H. Eckert, G. Schwering, Structure-Property Correlations in Lithium Phosphate Glasses: New Insights from 31 P ↔ 7 Li Double-Resonance NMR, Chem. Mater., 12 (2000) 1840–1846.

[38] L. van Wüllen, triple resonance transfer of populations in double resonance experiments for the detection of dipolar interactions, Solid State Nucl. Magn. Reson., 13 (1998) 123–127.

[39] M. Bak, J.T. Rasmussen, N.C. Nielsen, SIMPSON: a general simulation program for solid-state NMR spectroscopy, J. Magn. Reson., 147 (2000) 296–330.

[40] K. Takegoshi, S. Nakamura, T. Terao, C-13-H-1 dipolar-assisted rotational resonance in magic-angle spinning NMR, Chem. Phys. Lett., 344 (2001) 631–637.

[41] D. Massiot, F. Fayon, M. Capron, I. King, S. Le Calve, B. Alonso, J.O. Durand, B. Bujoli, Z.H. Gan, G. Hoatson, Modelling one- and two-dimensional solid-state NMR spectra, Magn. Reson. Chem., 40 (2002) 70–76.

[42] A. Samoson, Satellite transition high-resolution NMR of quadrupolar nuclei in powders, Chem. Phys. Lett., 119 (1985) 29–32.

[43] A.W. Hing, S. Vega, J. Schaefer, Transferred-Echo Double-Resonance NMR, J. Magn. Reson., 96 (1992) 205–209.

[44] T. Gullion, J. Schaefer, Rotational-Echo Double-Resonance NMR, J. Magn. Reson., 81 (1989) 196–200.

[45] A. Haimovich, A. Goldbourt, Characterization of lithium coordination sites with magic-angle spinning NMR, J. Magn. Reson., 254 (2015) 131–138.

[46] A. Haimovich, U. Eliav, A. Goldbourt, Determination of the Lithium Binding Site in Inositol Monophosphatase, the Putative Target for Lithium Therapy, by Magic-Angle-Spinning Solid-State NMR, J. Am. Chem. Soc., 134 (2012) 5647–5651.

[47] M. Bertmer, H. Eckert, Dephasing of spin echoes by multiple heteronuclear dipolar interactions in rotational echo double resonance NMR experiments, Solid State Nucl. Magn. Reson., (1999).

[48] U. Eliav, A. Haimovich, A. Goldbourt, Site-resolved multiple-quantum filtered correlations and distance measurements by magic-angle spinning NMR: Theory and applications to spins with weak to vanishing quadrupolar couplings, J. Chem. Phys., 144 (2016) 024201.

[49] X. Xue, M. Kanzaki, Proton Distributions and Hydrogen Bonding in Crystalline and Glassy Hydrous Silicates and Related Inorganic Materials: Insights from High-Resolution Solid-State Nuclear Magnetic Resonance Spectroscopy, J. Am. Ceram. Soc., 92 (2009) 2803–2830.

[50] M. Matthies, G. Zundel, Hydration and self-association of adenosine triphosphate, adenosine diphosphate, and their 1:1 complexes with magnesium(II) at various pH values: Infrared investigations, J. Chem. Soc. Perkin Trans. 2, (1977) 1824–1830.

[51] J.L. Oscarson, P. Wang, S.E. Gillespie, R.M. Izatt, G.D. Watt, C.D. Larsen, J.A.R. Renuncio, Thermodynamics of protonation of AMP, ADP, and ATP from 50 to 125°C, J. Solution Chem., 24 (1995) 171–200.

[52] R.N. Goldberg, Y.B. Tewari, Thermodynamics of the disproportionation of adenosine 5′-diphosphate to adenosine 5′-triphosphate and adenosine 5′-monophosphate I Equilibrium model, Biophys. Chem., 40 (1991) 241–261.

[53] R.M. Smith, A.E. Martell, Y. Chen, Critical Evaluation of Stability Constants for Nucleotide Complexes with Protons and Metal Ions and the Accompanying Enthalpy Changes, PURE Appl. Chem., 63 (1991) 1015–1080.

[54] A.L. Webber, S. Masiero, S. Pieraccini, J.C. Burley, A.S. Tatton, D. Iuga, T.N. Pham, G.P. Spada, S.P. Brown, Identifying guanosine self assembly at natural isotopic abundance by high-resolution 1H and 13C solid-state NMR spectroscopy, J. Am. Chem. Soc., 133 (2011) 19777–19795.

[55] M. Dracínský, P. Hodgkinson, Solid-state NMR studies of nucleic acid components, RSC Adv., 5 (2015) 12300–12310.

[56] E. Vinogradov, P.K. Madhu, S. Vega, Proton spectroscopy in solid state nuclear magnetic resonance with windowed phase modulated Lee–Goldburg decoupling sequences, Chem. Phys. Lett., 354 (2002) 193–202.

[57] M. Leskes, P.K. Madhu, S. Vega, Supercycled homonuclear dipolar decoupling in solid-state NMR: toward cleaner 1H spectrum and higher spinning rates, J. Chem. Phys., 128 (2008) 052309.

[58] E. Nimerovsky, R. Gupta, J. Yehl, M. Li, T. Polenova, A. Goldbourt, Phase-modulated LA-REDOR: a robust, accurate and efficient solid-state NMR technique for distance measurements between a spin-1/2 and a quadrupole spin, J. Magn. Reson., 244 (2014) 107–13.

[59] M. Makrinich, E. Nimerovsky, A. Goldbourt, Pushing the limit of NMR-based distance measurements – retrieving dipolar couplings to spins with extensively large quadrupolar frequencies, Solid State Nucl. Magn. Reson., 92 (2018) 19–24.

[60] R.D. Shannon, Revised effective ionic radii and systematic studies of interatomic distances in halides and chalcogenides, Acta Crystallogr. Sect. A, 32 (1976) 751–767.

