## supplementary information for "How does the mood stabilizer lithium bind ATP, the energy currency of the cell"

#### NMR Experimental Parameters

##### $^{31}P$ CPMAS NMR of Li-ATP (FIGURE 1)

The spectrum was recorded at a field of 14.1 T, a spinning rate of 14 kHz and a temperature set to 0 °C. It consists of 4 scans with a recycle delay of 15 s. CP time was 1.5 ms. $^{1}H$ decoupling was performed using the $SW_f$-TPPM technique with a radio-frequency (rf) power of 80 kHz. The spectrum was processed with line broadening of 20 Hz.

##### $^{31}P$-$^{31}P$ DARR on {23%-$^{7}Li$}-Li-ATP (FIGURE 2)

| sequence / parameters | DARR |
|---|---|
| $^{1}H$ frequency [MHz] | 599.98 |
| Spinning frequency ($\nu_r$ ) [kHz] | 14 |
| Set temperature [°C] | 0 |
| Acquisition points ($t_1$/$t_2$) | 140/4990 |
| Acquisition times ($t_1$/$t_2$) [ms] | 10/25 |
| Carrier frequency [ppm] | -12 |
| Pulse power level ($\nu_P$) [kHz] | 93 |
| CP power level ($\nu_H$/$\nu_P$) [kHz] | ~58 (ramp 90-100%)/~50 |
| $^{1}H$-$^{31}P$ CP contact time [ms] | 1500 |
| DARR mixing time [ms] | 15, 100 |
| $^{1}H$ Decoupling [kHz]<br>$Sw_f$-TPPM decoupling (tangent pulse, 78-122%) | 80 |
| Relaxation delay [sec] / Scans | 15.0/16 |
| Spectral width (F1/F2) [kHz] | 7/100 |
| **processing parameters F1/F2** | |
| Processing software | TopSpin |
| Total # of points (F1/F2) | 512/16384 |
| Apodization functions | 1. Lorentz to Gauss transformation in F1: (GmaxPos0.1/ WidthHz(-20));<br>2. Exponential in in F2: 20Hz |

##### $^{31}P$ CPMAS NMR of $Na_2ATP$ and Li-ATP (FIGURE 3)

Both spectra were recorded at a field of 9.4 T, a spinning rate of 14 kHz and a temperature set to -20 °C. They consisted of 4 scans, with recycle delays of 11 s and 8 s for Na and Li ATP experiments, respectively, and CP contact times of 1.5 ms. $^{1}H$

decoupling was performed using the $SW_f$-TPPM technique with rf power of 80 kHz. The spectra were processed with line broadening of 20 Hz.

#### 2D $^7$Li-$^{31}$P TEDOR on {23%-$^7$Li}-Li-ATP (Figure 4)

| sequence / parameters | TEDOR |
|---|---|
| $^1$H frequency [MHz] | 599.98 |
| Spinning frequency ($\nu_r$ ) [kHz] | 14 |
| Set temperature [°C] | 0 |
| Acquisition points ($t_1/t_2$) | 100/3988 |
| Acquisition times ($t_1/t_2$) [ms] | 7/20 |
| Carrier frequency ($^{31}$P/$^7$Li) [ppm] | -13.6/0 |
| Pulse power level ($\nu_P/\nu_{Li}$) [kHz] | 50/50 |
| CP power level ($\nu_H/\nu_P$) [kHz] | ~58 (ramp 90-100%)/~50 |
| $^1$H-$^{31}$P CP contact time [ms] | 2000 |
| Preparation period *n* [×$t_r$] ([ms]) | 10 (714ms) |
| Reconversion period *m* [×$t_r$] | 18 |
| $^1$H Decoupling [kHz]<br>$Sw_f$-TPPM decoupling (tangent pulse, 78-122%) | 80 |
| Relaxation delay[sec] / Scans | 12.0/16 |
| Spectral width (F1/F2) [kHz] | 7/100 |
| **processing parameters F1/F2** | |
| Processing software | TopSpin |
| Total # of points (F1/F2) | 512/16384 |
| Apodization functions | Exponential in (F1;F2): (50Hz; 70Hz) |

#### $^{31}$P{$^7$Li} REDOR on {~23%-$^7$Li}-Li-ATP (Figures 5-6)

The spectra were recorded at a field of 14.1 T, a spinning rate of 14 kHz and a temperature set to 0 °C. It consisted of 32 scans with a recycle delay of 12 s, CP time of 2 ms and $SW_f$-TPPM decoupling (80 kHz). REDOR dephasing times were taken at 2-62 rotor periods ($T_R$), with increments of 2 up to 26 $T_R$ and increments of 4 up to 62 $T_R$. The $^7$Li $\pi$ pulses followed the XY8 phase cycling scheme. Figure 5 presents raw data and in figure 6 the data is deconvoluted.

#### *w*PMLG-HETCOR on {23%-$^7$Li}-Li-ATP (Figures 7, 8)

| sequence / parameters | *w*PMLG -HETCOR | | |
|---|---|---|---|
| | **$^1$H-$^{31}$P** | **$^1$H-$^{31}$P** | **$^1$H-$^7$Li** |
| $^1$H frequency [MHz] | 599.98 | | |
| Spinning frequency ($\nu_r$ ) [kHz] | 11.5 | | |
| Set temperature [°C] | 0 | | -20 |
| Acquisition points ($t_1/t_2$) | 120/4990 | 220/4990 | 120/4990 |
| Acquisition times ($t_1/t_2$) [ms] | 3/25 | 6/25 | 3/25 |
| Carrier frequency ($^1$H/X) [ppm] | 8.7$^a$ / -12.7 | | 8.7$^a$ / 0 |

| | | | |
|---|---|---|---|
| $^1H$ pulse power level ($\nu_H$) [kHz] | 94.3 | | |
| CP power level ($\nu_H/\nu_X$) [kHz] | ~55 (ramp 90-100%)/~50 | | ~67 (ramp 90-100%)/~50 |
| $^1H$-X CP contact time [ms] | 200, 500, 1000 | 1000, 3000, 7000 | 500 |
| PMLG pulse [ms] | 1.2 | | |
| PMLG window [ms] | 6 | | |
| $^1H$ Decoupling [kHz]<br>$Sw_f$-TPPM decoupling (tangent pulse, 78-122%) | 80 | | |
| Relaxation delay [sec] / Scans | 15.0/8 | | 12.0/16 |
| Spectral width (F1/F2) [kHz] | 18.5 (3 PMLG cycles)/100 | | |
| **processing parameters F1/F2** | | | |
| Processing software | TopSpin | | |
| Total # of points (F1/F2) | 512/16384 | | 256/16384 |
| Apodization functions | Exponential in (F1;F2): (10Hz; 10Hz) | | Exponential in (F1;F2): (10Hz; 30Hz) |

[a] Measured for a simple SPE experiment

#### X-$^1$H HETCOR on {NA $^7$Li}-Li-ATP (Figures 7c, 8b)

| sequence / parameters | **Proton detected HETCOR** | |
|---|---|---|
| | **$^{31}P$-$^1H$** | **$^7Li$-$^1H$** |
| $^1H$ frequency [MHz] | 599.85 | |
| Spinning frequency ($\nu_r$) [kHz] | 62 | |
| Set temperature [°C] | -30 | |
| Acquisition points ($t_1/t_2$) | 186/1988 | 92/1988 |
| Acquisition times ($t_1/t_2$) [ms] | 15/10 | |
| Carrier frequency ($^1H$/X) [ppm] | 8.5[a] / -13.6 | 8.4/ -0.1 |
| $\pi/2$ pulse power level ($\nu_H/\nu_X$) [kHz] | 150/100 | |
| CP power level ($\nu_H/\nu_X$) [kHz]<br>Ramp: $^1$HX 90-100% / X$^1$H 100-90% | ~70 /~20 | |
| CP contact times [ms] | 2750 | 1700 |
| $^1H$ Decoupling [kHz]<br>$Sw_f$-TPPM decoupling (tangent pulse, 78-122%) | 15 | |
| X Decoupling [kHz]<br>WALTZ-16 decoupling | 10 | |
| Relaxation delay [sec] / Scans | 12.0/16 | |
| Spectral width (F1/F2) [kHz] | 6.2/100 | 3.1/100 |
| **processing parameters F1/F2** | | |
| Processing software | TopSpin | |
| Total # of points (F1/F2) | 512/8192 | 256/8192 |
| Apodization functions | 1. Exponential in F1: 10Hz<br>2. Squared sine-bell in F2: shifted by 90° | 1. Exponential in F1: 30Hz<br>2. Squared sine-bell in F2: shifted by 90° |

##### $^{23}Na$ spectra of $Na_2ATP$ and Li-ATP (Figure 9)

The spectra were recorded at a field of 9.4 T, a spinning speed of 14 kHz, at T = -20 °C. Both were acquired with a single scan and processed with line broadening of 50 Hz.

##### $^{7}Li\{^{23}Na\}$ PM-RESPDOR distance measurement experiments (Figure 10a)

The spectra were recorded at a field of 14.1 T, a spinning speed of 14 kHz, T = -15°C, using 32, 64 or 96 scans with a recycle delay of 19.65 s, The observed $^{7}Li$ nucleus was excited via CP time of 1 ms and $SW_f$-TPPM decoupling (80 kHz) was applied. Dephasing times were taken at 2-170 $T_R$, with increments of 8 up to 98 $T_R$, increments of 16 up to 144 $T_R$, and a final increment of 24. The $^{7}Li$ $\pi$ pulses followed the XY64 phase cycling scheme. The phase-modulated pulse on $^{23}Na$ lasted 0.714 ms ($10T_R$). The pulse shape was given elsewhere.[1]

##### $^{31}P\{^{23}Na\}$ PM-RESPDOR distance measurement experiments (Figure 10b)

Experimental parameters appear in Figure S7.

##### $^{23}Na\{^{7}Li\}$ Central-transition REDOR experiments (Figure 11)

The spectra were recorded at a field of 14.1 T, a spinning speed of 14 kHz, T = -15 °C, using 64 and 128 scans with a recycle delay of 5 s. The observed $^{23}Na$ nucleus was excited via selective central-transition excitation. Dephasing times were taken at 34-322 $T_R$. The $^{7}Li$ $\pi$ pulses followed the XY8 phase cycling scheme.

### Figures and Tables

#### $^{31}P$-$^{31}P$ DARR on $Na_2ATP$

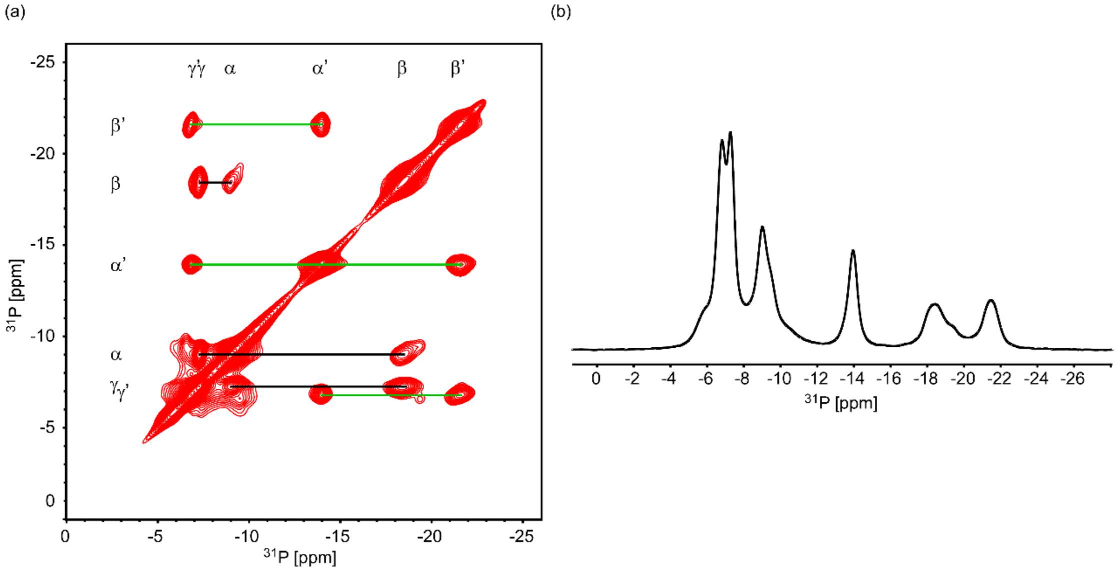


**Figure S1.** Spectra of disodium ATP trihydrate recrystallized from a commercial material. (a) 2D $^{31}P$-$^{31}P$ DARR of $Na_2ATP$ acquired with a short mixing time of 15 ms. The lines indicate correlations between adjacent phosphate groups, black lines correlating signals within phosphates of one ATP molecule ($\alpha$, $\beta$, $\gamma$), and the green lines for the second ATP molecule ($\alpha'$, $\beta'$, $\gamma'$). It can be seen that the position of the gamma phosphates is reversed in comparison with Li-ATP. (b) The 1D $^{31}P$ CPMAS spectrum of $Na_2ATP$.

#### Simulations of $^{31}P\{^{7}Li\}$ REDOR

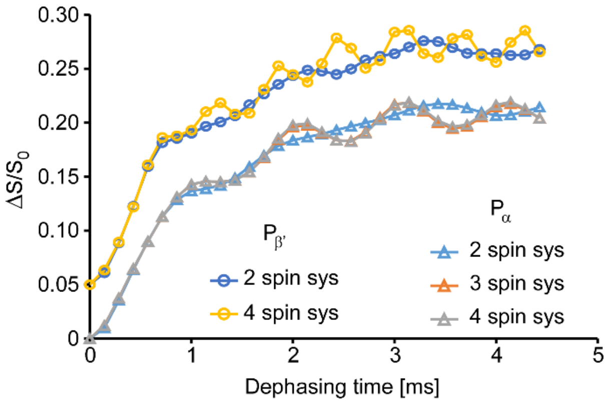


**Figure S2**. SIMPSON simulations of $^{31}P\{^{7}Li\}$ REDOR for different spin system sizes. Detection is on a single phosphorus atom. We simulated two spin systems of Li-$P_{\alpha/\beta'}$, a three spin system of Li-$P_\alpha$-$P_\beta$ and a four spin system of Li-$P_\alpha$-$P_\beta$-$P_{\beta'}$, taking into account isotropic shifts, CSA, Cq and all homonuclear and heteronuclear interactions. The initial rise of the REDOR curves is unaffected by the spin system size. The simulated curves for $P_{\beta'}$ were shifted by 0.05 $\Delta S/S_0$ units for clarity. The simulations take into account the 23% labeling of lithium. Thus, the maximum value of the $\Delta S/S_0$

curve is 0.23 (with oscillations around it). The ‘initial rise’ is therefore around 65% of the maximal possible recoupling.

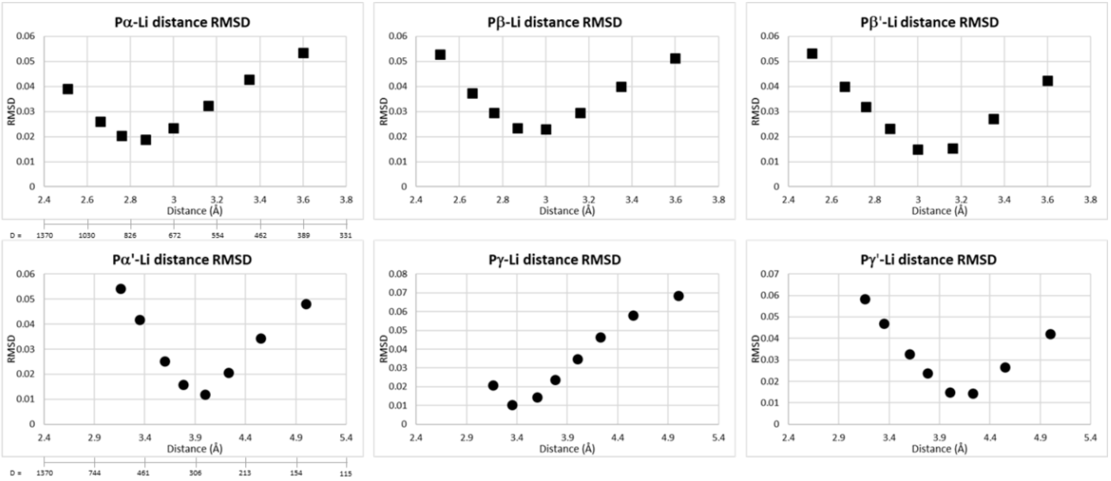


**Figure S3**. RMSD plots generated for all six $^{31}P\{^{7}Li\}$ REDOR curves shown in figure 6. Top row is for the closest phosphate species. The dipolar coupling constants (in Hz) corresponding to the distances shown on the x-axis are indicated on the left column. The following number of points on the curves were used in the RMSD calculations: Pα 4 pts, Pβ 4 pts, Pβ’ 5 pts, Pα’ 11 pts, Pγ 9 pts, Pγ’ 13 pts.

#### $^{1}$H Solution NMR for LiATP

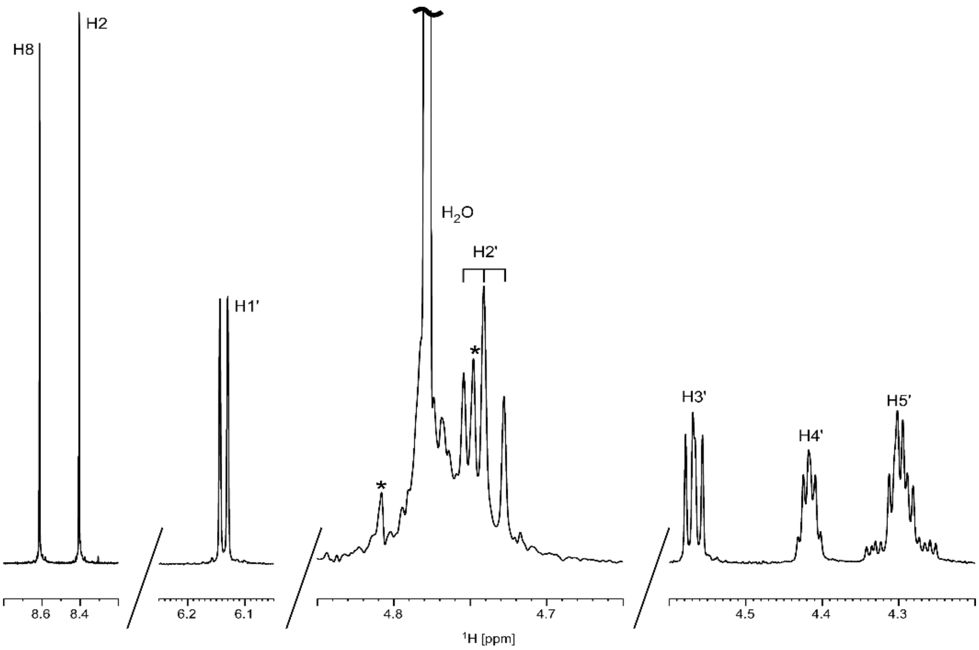


**Figure S4.** $^{1}$H solution NMR spectrum of Li-ATP in $D_2O$ taken in a 9.4 T spectrometer. The spectrum is referenced to acetone at 2.2 ppm. Spinning side bands are marked with asterisks. The water signal is truncated for clarity and its spinning side bands are marked with asterisks.

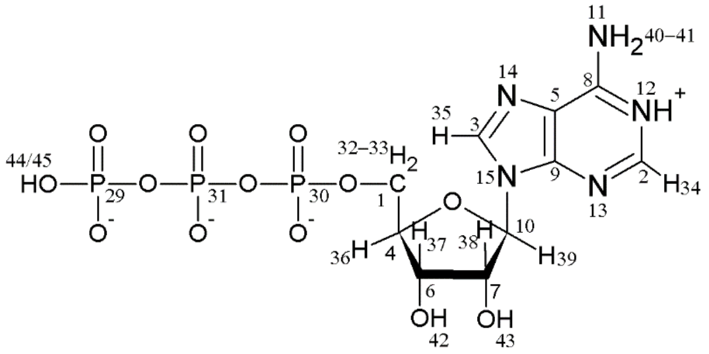


**Scheme S1:** Atom numbering of ATP according to BMRB entries bmse000006, bmse000854, and bmse000993.

#### Table S1. Assignment of $^{1}$H NMR shifts in Li-ATP

| Solution shift (ppm) | Assignment | Assignment, BMRB format | Solid state shift (ppm) | Assignment | Remarks |
|---|---|---|---|---|---|
| | | | 15.3 | N1H$^{+}$ | Correlates P$\gamma$/$\gamma$'. Correlates lithium only at long mixing times. |
| | | | 11.6 | P$\gamma$/$\gamma$'-OH | Fast P$\gamma$/$\gamma$' build-up. Correlates lithium only at long mixing times |
| | | | 10.0 | | Appears at long mixing times at fast MAS. Correlates to P$_{\beta'}$ (in P-H) and to lithium (in Li-H) |
| 8.6 | H8 | H35 | 8.7 | H2/H8 (ATP) | Correlates with $\alpha$', $\beta$', $\gamma$' in HETCOR 1 ms |
| 8.4 | H2 | H34 | 8.0 | H2/H8 (ATP) | Correlates with $\alpha$, $\beta$, $\gamma$ in HETCOR 1 ms*. Correlates lithium stronger then 8.7. |
| 6.15 | H1' | H39 | 7.0 | H1' | Fast build-up with P$_{\alpha}$. |
| 4.78 | H2O | | ~5 ppm | $H_2O$ or H10 ($NH_2$) | Broad signal |
| 4.75 | H2' | H43 | 4.9 | H2'-H5' | Fast build-up with P$_{\alpha}$. Slower for P$_{\alpha'}$. Weaker with P$_{\beta/\beta'/\gamma/\gamma'}$. Correlates lithium at short mixing times. |
| 4.58 | H3' | H42 | 4.2 | H2'-H5' | Fast build-up with P$_{\alpha}$. Slower for P$_{\alpha'}$. Weaker with P$_{\beta/\beta'/\gamma/\gamma'}$. |
| 4.42 | H4' | H36 | | | |
| 4.3 | H5' | H32-33 | | | |

* Alternatively, that the two peaks represent each a different base hydrogen and the spatial arrangement of the two ATP molecules can render each hydrogen closer to a different phosphate chain.

#### Table S2. pKa values of ATP

| Atom | pKa | Reference |
|---|---|---|
| Imino N1 | 3.98-4.96; 25 °C | [2–5] |
| Ribose OH | ~12-13 | [2,6] |
| OH of P$\alpha$, P$\beta$, P$\gamma$ | 1-2.17; 25 °C | [2–5] |
| Second OH of P$\gamma$ | ~6.48-7.68; 25 °C | [2–5] |

#### $^{1}H$-$^{31}P$ CP buildup curves

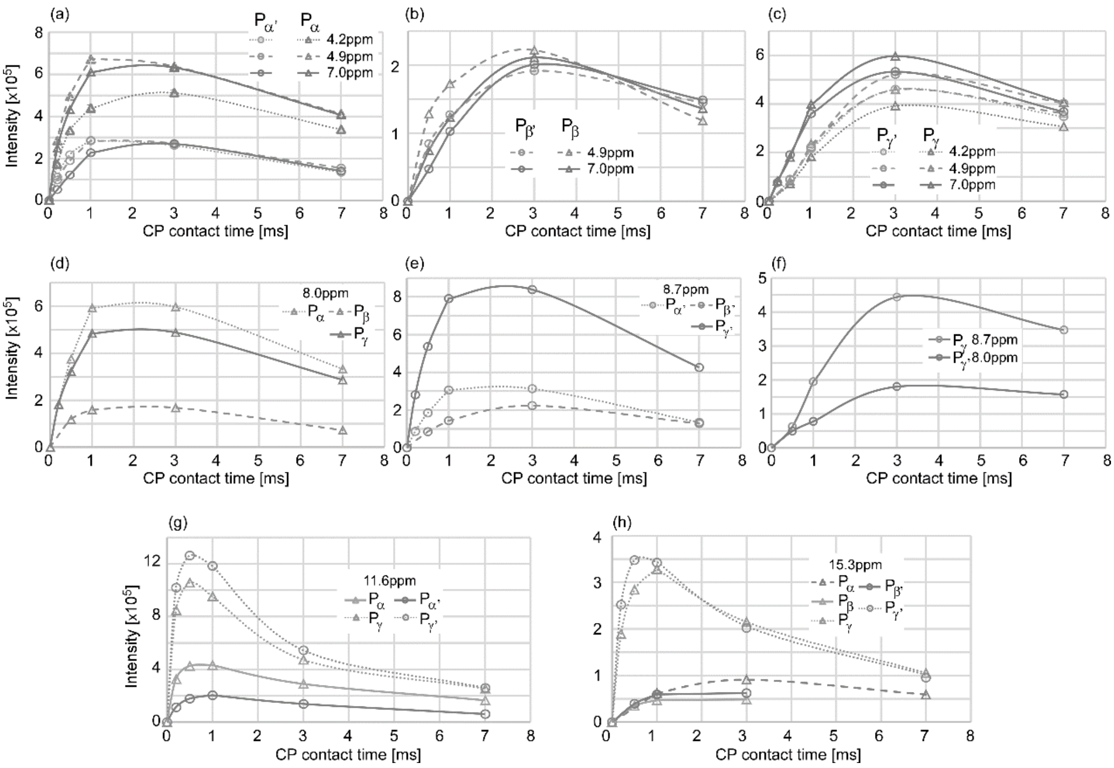


**Figure S5.** CP buildup curves of $\boldsymbol{wPMLG_{m\bar{m}}^{x\bar{x}}}$ $^{1}H$-$^{31}P$ crosspeaks. Two different datasets were used and the data points were factored according to a CP contact time of 1 ms existing in both datasets. Buildup curves for the correlations with $^{1}H$ peaks at 4.2, 4.9 and 7.0 ppm are presented in (a) for $P_{\alpha}$ and $P_{\alpha'}$, (b) for $P_{\beta}$ and $P_{\beta'}$ and (c) for $P_{\gamma}$ and $P_{\gamma'}$. Buildup curves are presented for the correlations with $^{1}H$ peaks at (d) and (f) 8.0 ppm, (e) and (f) 8.7 ppm, (g) 11.6 ppm and (h) 15.3 ppm.

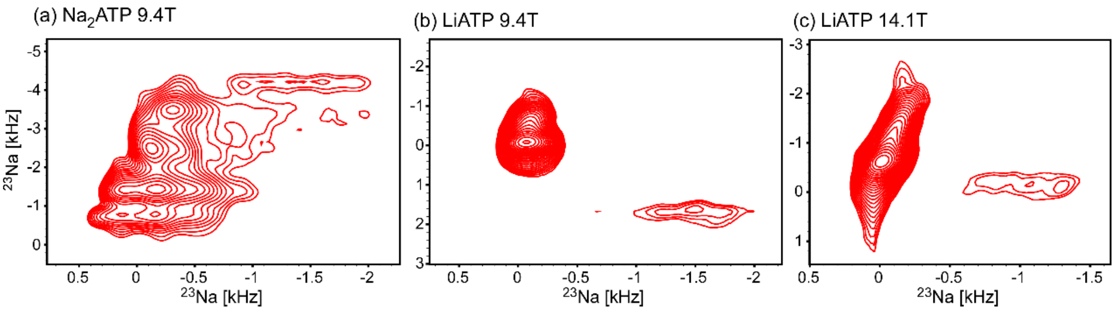


**Figure S6.** $^{23}Na$ MQMAS spectra of (a) recrystallized disodium ATP at 9.4 T; (b) {24%-$^{7}Li$}-Li-ATP at 9.4T; (c) {24%-$^{7}Li$}- $^{7}Li$-ATP at 14.1 T.

##### $^{31}$P{$^{23}$Na} PM-RESPDOR distance measurement experiments

The spectra were recorded at a field of 14.1 T, at different spinning speeds of 12, 14 and 14.1 kHz, T = -15 °C, using 32 scans with a recycle delay of 19.65 s. The observed $^{31}$P nucleus was excited via a CP time of 1.5 ms and SW$_f$-TPPM $^1$H decoupling (80 kHz) was applied. Dephasing times were taken at 2-42 rotor periods ($T_R$). The $^{31}$P $\pi$ pulses followed the XY64 phase cycling scheme. The phase-modulated (PM) pulse on $^{23}$Na lasted 0.833, 0.714 and 0.709 ms (10$T_R$) for all spinning speeds and for 14.1 kHz data was also measured with a PM pulse 1.418 ms (20$T_R$). The pulse shape is given elsewhere.[1]

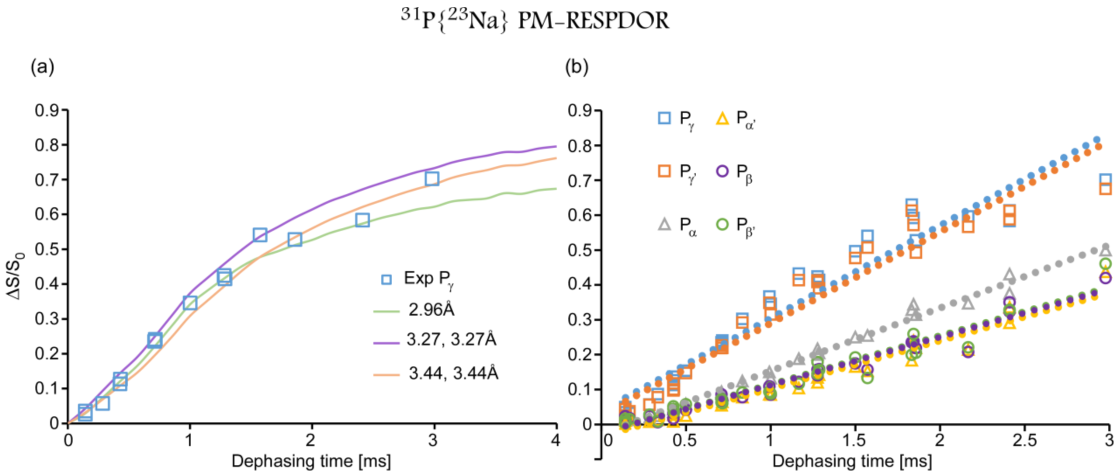


**Figure S7**. (a) $^{31}$P{$^{23}$Na} PM-RESPDOR recoupling curve of the $P_\gamma$ phosphate of Li-ATP measured at MAS of 14 & 14.1 kHz with a phase-modulated pulse of 10$T_R$. The experimental points $\Delta S/S_0$ are shown as squares. A fit to an isolated pair yields a phosphate-sodium distance of ~3Å, which is a short distance in comparison to reported Na-P distances in sodium ATP. SIMPSON simulated curves for two sodium sites present a better fit for the experimental data. Presented are simulations for equal distances of 3.27 and 3.44Å. (b) PM-RESPDOR $^{31}$P{$^{23}$Na} recoupling curves for all six phosphates measured at the experimental conditions depicted above. Linear trend-lines were drawn in order to portray the differences in the slopes of the recoupling curves indicating on the different inter-nuclear distances.

##### $^{23}$Na{$^{31}$P} REDOR measurements

The spectra were recorded at a field of 14.1 T, a spinning speed of 14.1 kHz, T = -15 °C, using 16, 32 and 64 scans with a recycle delay of 5 s. The observed $^{23}$Na nucleus was excited via a selective central-transition excitation. Dephasing times were taken at 6-98 $T_R$. The $^{31}$P $\pi$ pulses followed the XY8 phase cycling scheme.

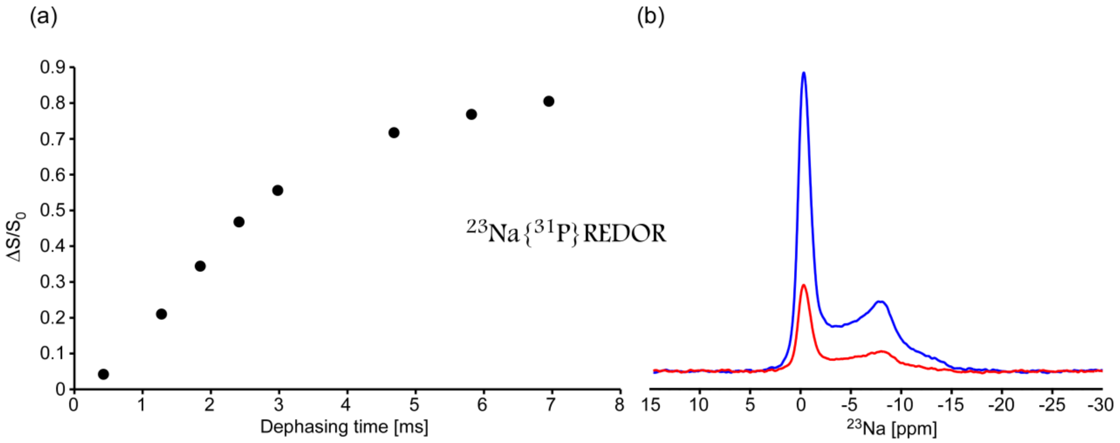


**Figure S8.** (a) $^{23}$Na{$^{31}$P} REDOR data points represent the integral of the entire sodium signal. (b) A single $\Delta S/S_0$ point acquired at a mixing times of 4.68 ms showing the $S_0$ reference signal (blue) and the decay signal S (red). It can be seen that the dephasing seems to be uniform.

#### Crystallographic models of metal-ATP complexes.

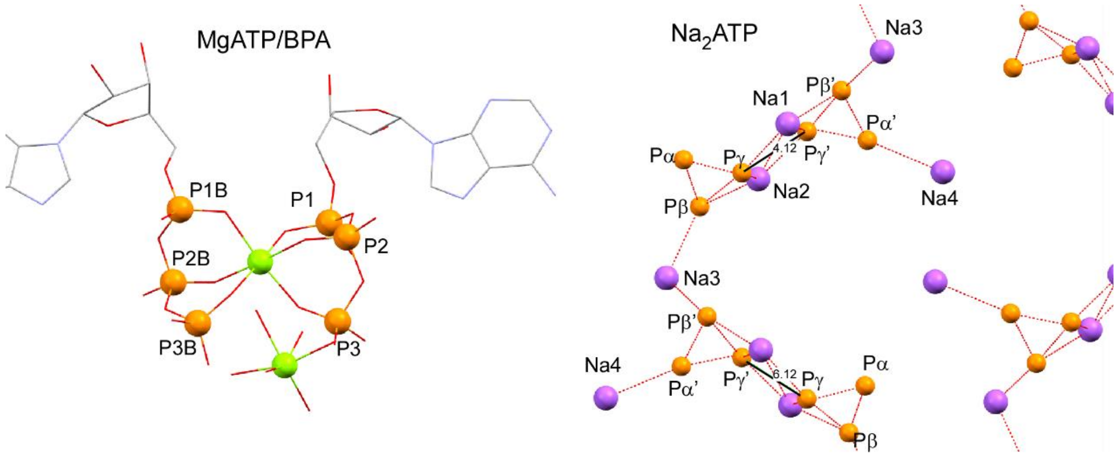


**Figure S9.** Crystallographic structures of (left) MgATP/BPA ([Mg(H20)$_6$[H- bis(2-pyridyl)amine]$_2$[Mg(Hatp)$_2$]•12$H_2O$), CCDC identifier DECDIY, Cini et al.[7]. $Mg^{2+}$ ions are in green, phosphorous atoms in orange. The BPA molecule is dropped for clarity and the two ATP units are symmetrically equivalent. CaATP/BPA has an almost identical structure. (right) $Na_2$ATP•3$H_2O$, CCDC identifier ADENTP, Kennard et al. [8]. The plots depict a part of the unit cell demonstrating the positions of the four sodium ions (violet) and the relative positions of the phosphate moieties in the dimer. One of the dimers has a Pγ-Pγ' distance of 4.12Å, the other of 6.12Å. This is in agreement with the NMR observations of Potrzebowski et al.[9] but not with our Li-ATP model. The adenosine moiety has been omitted for clarity. Plots were generated using Mercury version 3.10.3 and rendered using POV-ray.

### NMR Experimental Parameters, 2D experiments

#### $^{31}P$-$^{31}P$ DARR on $Na_2ATP$ (FIGURE S1)

| sequence / parameters | DARR |
|---|---|
| $^{1}H$ frequency [MHz] | 400.2 |
| Spinning frequency ($\nu_r$ ) [kHz] | 14 |
| Set temperature [°C] | -20 |
| Acquisition points ($t_1/t_2$) | 238/4990 |
| Acquisition times ($t_1/t_2$) [ms] | 17/25 |
| Carrier frequency [ppm] | -14.9 |
| Pulse power level ($\nu_P$) [kHz] | 50 |
| CP power level ($\nu_H/\nu_P$) [kHz] | ~65 (ramp 90-100%)/~50 |
| $^{1}H$-$^{31}P$ CP contact time [ms] | 1500 |
| DARR mixing time [ms] | 15 |
| $^{1}H$ Decoupling [kHz]<br>$SW_f$-TPPM decoupling (tangent pulse, 78-122%) | ~80 |
| Relaxation delay [sec] / Scans | 8.0/16 |
| Spectral width (F1/F2) [kHz] | 7/100 |
| **processing parameters F1/F2** | |
| Processing software | TopSpin |
| Total # of points (F1/F2) | 1024/16384 |
| Apodization functions | 1. Lorentz to Gauss transformation in F1: (GmaxPos0.1/ WidthHz(-20));<br>2. Exponential in in F2: 20Hz |

#### $^{23}Na$ MQMAS on $Na_2ATP$ (FIGURE S6)

| sequence / parameters | MQMAS |
|---|---|
| $^{1}H$ frequency [MHz] | 400.2 |
| Spinning frequency ($\nu_r$ ) [kHz] | 14 |
| Set temperature [°C] | -20 |
| Acquisition points ($t_1/t_2$) | 160/1598 |
| Acquisition times ($t_1/t_2$) [ms] | 2/8 |
| Carrier frequency [ppm] | 11.7 |
| Selective CT Pulse power level ($\nu_P$) [kHz] | 14.7 |
| Excitation pulse<br>length ($\mu$s) / power level (kHz) | 6.8 / 198 |
| FAM pulses<br>length ($\mu$s) / power level (kHz) | 2 / 100 |
| FAM cycles | 5 |
| Relaxation delay [sec] / Scans | 0.13/1440 |
| Spectral width (F1/F2) [kHz] | 40/100 |

| processing parameters F1/F2 | |
|---|---|
| Processing software | TopSpin |
| Total # of points (F1/F2) | 512/4096 |
| Apodization functions | Squared sine-bell in (F1;F2): shifted by (90°; 0°) |

#### $^{23}$Na MQMAS on LiATP (FIGURE S6)

| sequence / parameters | MQMAS | |
|---|---|---|
| $^{1}$H frequency [MHz] | 400.2 | 599.85 |
| Spinning frequency ($\nu_r$ ) [kHz] | 14 | 14 |
| Set temperature [°C] | -20 | -15 |
| Acquisition points ($t_1/t_2$) | 80/2390 | 80/2988 |
| Acquisition times ($t_1/t_2$) [ms] | 2.2/12 | 2.2/15 |
| Carrier frequency [ppm] | 2.5 | 7.4 |
| Selective CT Pulse power level ($\nu_P$) [kHz] | 30 | 15 |
| Excitation pulse length ($\mu$s) / power level (kHz) | 6.6 / 160 | 8.5 / 90 |
| FAM pulses length ($\mu$s) / power level (kHz) | 1.5 / 100 | 2 / 95 |
| FAM cycles | 5 | 5 |
| Relaxation delay [sec] / Scans | 0.66/96 | 1.2/240 |
| Spectral width (F1/F2) [kHz] | 18/100 | 18/100 |
| **processing parameters F1/F2** | | |
| Processing software | TopSpin | TopSpin |
| Total # of points (F1/F2) | 256/8192 | 256/8192 |
| Apodization functions | Squared sine-bell in (F1;F2): shifted by (90°; 0°) | Squared sine-bell in (F1;F2): shifted by (90°; 0°) |

#### References


[1] E. Nimerovsky, R. Gupta, J. Yehl, M. Li, T. Polenova, A. Goldbourt, *J. Magn. Reson.* **2014**, *244*, 107–13.

[2] M. Matthies, G. Zundel, *J. Chem. Soc. Perkin Trans. 2* **1977**, 1824–1830.

[3] J. L. Oscarson, P. Wang, S. E. Gillespie, R. M. Izatt, G. D. Watt, C. D. Larsen, J. A. R. Renuncio, *J. Solution Chem.* **1995**, *24*, 171–200.

[4] R. N. Goldberg, Y. B. Tewari, *Biophys. Chem.* **1991**, *40*, 241–261.

[5] R. M. Smith, A. E. Martell, Y. Chen, *PURE Appl. Chem.* **1991**, *63*, 1015–1080.

[6] H. Aström, E. Limén, R. Strömberg, *J. Am. Chem. Soc.* **2004**, *126*, 14710–1.

[7] R. Cini, M. C. Burla, A. Nunzi, G. P. Polidori, P. F. Zanazzi, *J. Chem. Soc., Dalt. Trans.* **1984**, 2467–2476.

[8] O. Kennard, N. W. Isaacs, W. D. S. Motherwell, J. C. Coppola, D. L. Wampler, A. C. Larson, D. G. Watson, *Proc. R. Soc. A Math. Phys. Eng. Sci.* **1971**, *325*, 401–436.

[9] M. J. Potrzebowski, J. Gajda, W. Ciesielski, I. M. Montesinos, *J. Magn. Reson.* **2006**, *179*, 173–181.
